# Global distribution of anaerobic dichloromethane degradation potential

**DOI:** 10.1101/2021.08.30.458270

**Authors:** Robert W. Murdoch, Gao Chen, Fadime Kara Murdoch, E. Erin Mack, Manuel I. Villalobos Solis, Robert L. Hettich, Frank E. Löffler

**Affiliations:** Center for Environmental Biotechnology, University of Tennessee, Knoxville, TN 37996, USA; Department of Civil and Environmental Engineering, University of Tennessee, Knoxville, TN 37996, USA; Department of Microbiology, University of Tennessee, Knoxville, TN 37996, USA; Department of Biosystmes Engineering and Soil Science, University of Tennessee, Knoxville, TN 37996, USA; Corteva Environmental Remediation, Corteva Agriscience, Wilmington, DE 19805, USA; Biosciences Division, Oak Ridge National Laboratory, Oak Ridge, TN 37831, USA

**Keywords:** Dichloromethane fluxes, Degradation Potential, Bioremediation, Ozone Destruction

## Abstract

Anthropogenic activities and natural processes release dichloromethane (DCM), a toxic chemical with substantial ozone-depleting capacity. Specialized anaerobic bacteria metabolize DCM; however, the genetic basis for this process has remained elusive. Comparative genomics of the three known anaerobic DCM-degrading bacterial species revealed a homologous gene cluster, designated the methylene chloride catabolism (*mec*) gene cassette, comprising eight to ten genes with predicted 79.6 – 99.7% amino acid identity. Functional annotation identified genes encoding a corrinoid-dependent methyltransferase system, and shotgun proteomics applied to two DCM-catabolizing cultures revealed high expression of proteins encoded on the *mec* gene cluster during anaerobic growth with DCM. In a DCM-contaminated groundwater plume, the abundance of *mec* genes strongly correlated with DCM concentrations (R^2^ = 0.71 – 0.85) indicating their value as process-specific bioremediation biomarkers. *mec* gene clusters were identified in metagenomes representing peat bogs, the deep subsurface, and marine ecosystems including oxygen minimum zones (OMZs), suggesting DCM turnover in diverse habitats. The broad distribution of anaerobic DCM catabolic potential suggests a relevant control function for emissions to the atmosphere, and a role for DCM as a microbial energy source in critical zone environments. The findings imply that the global DCM flux might be far greater than emission measurements suggest.

**Importance:** Dichloromethane (DCM) is an increasing threat to stratospheric ozone with both anthropogenic and natural emission sources. Anaerobic bacterial metabolism of DCM has not yet been taken into consideration as a factor in the global DCM cycle. The discovery of the *mec* gene cassette associated with anaerobic bacterial DCM metabolism and its widespread distribution in environmental systems highlight a strong attenuation potential for DCM. Knowledge of the *mec* cassette offers new opportunities to delineate DCM sources, enables more robust estimates of DCM fluxes, supports refined DCM emission modeling and simulation of the stratospheric ozone layer, reveals a novel, ubiquitous C_1_ carbon metabolic system, and provides prognostic and diagnostic tools supporting bioremediation of groundwater aquifers impacted by DCM.

## Introduction

Dichloromethane (DCM, methylene chloride) is a widely distributed halomethane, produced both naturally and industrially. While anthropogenic DCM has received attention due to widespread groundwater contamination and, more recently ozone destruction potential, analysis of Antarctic ice cores has demonstrated that DCM was present in the atmosphere prior to the industrial era at approximately 10% of modern levels (Trudinger et al., 2004). The natural sources of DCM are diverse, encompassing both abiotic (Isidorov et al., 1990; Kanters & Louw, 1996) and biotic (Eustáquio et al., 2008; Hoekstra et al., 1998; Wuosmaa & Hager, 1990) processes and are estimated to contribute up to one third of total emissions (Gribble, 2010). Since the 1960’s, atmospheric DCM concentrations rose steadily, with a mean annual increase of approximately 8% (Hossaini et al., 2017) although the reported worldwide production and use has been steady or declining since 2010 (McCulloch, 2017). Possible explanations include undocumented production, rogue emmissions, or increased natural emissions reflecting environmental (e.g., climate) change responses.

Atmospheric measurements and corresponding efforts to extrapolate to global-scale emissions have led to the perception that marine systems and biomass combustion (e.g., wildfires) are the primary non-industrial sources of DCM (Gribble, 2010), releasing estimated amounts of 190 and 60 Gg of DCM each year, respectively. Natural and deliberate forest fires have increased in frequency and size (Haines et al., 2020), a global trend that can be expected to lead to further formation of DCM. Wetlands emit up to 2 Gg DCM per year (Cox et al., 2004; Hu et al., 2017; Kolusu et al., 2018) (Supplementary Information), and volcanic activity contributes an estimated amount of 0.021 Gg/y (Gribble, 2010). Halomethanes occur in crustal minerals, and DCM release from rocks from the near-surface and deep subsurface have been reported (Mulder et al., 2013; Svensen et al., 2009). Knowledge gaps remain and not-yet identified environmental sources of DCM are likely. The current understanding of global DCM fluxes, as opposed to emissions, is very limited (Figure 1) (McCulloch, 2017).

**Figure 1.**
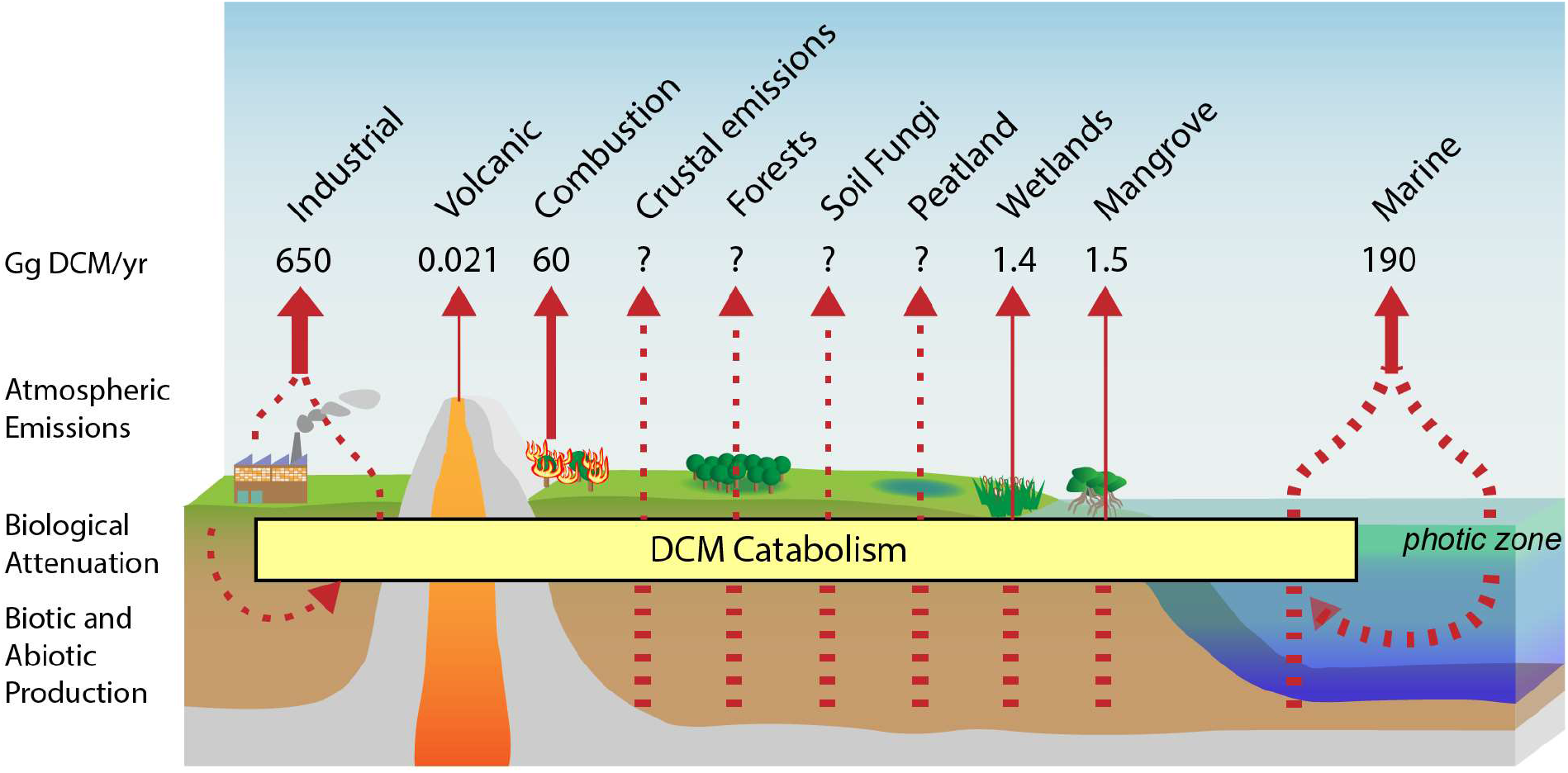
Major reported or potential DCM atmospheric emission sources. Width of the solid arrows is log-proportional to the magnitude of DCM emission estimates or potential. Dashed lines represent putative DCM sources, fluxes, and emissions that have not been directly investigated.

The rates of DCM production and consumption are unclear, and biological attenuation in anaerobic systems prior to release to the atmosphere has not been incorporated into existing atmospheric emission models (Hossaini et al., 2017).

Anaerobic metabolism of DCM was described in a bacterial isolate, *Dehalobacterium formicoaceticum* strain DMC (Defo) (*19*). Two additional bacterial populations responsible for DCM-metabolism in anaerobic enrichment cultures have been identified, ‘*Ca*. Dichloromethanomonas elyunquensis’ (Diel) (Justicia-Leon et al., 2012; Kleindienst et al., 2016, 2017) and ‘*Ca*. Formimonas warabiya’ (Dcmf) (Holland et al., 2021). All three DCM degraders are members of the family Peptococcaceae within the phylum Firmicutes. Bacterial dechlorination of DCM under oxic conditions is catalyzed by the glutathione-S-transferase (GST) DcmA (Muller et al., 2011); however, GST enzymes are generally absent in obligate anaerobes (Allocati et al., 2009). Accordingly, anaerobes metabolize DCM following a distinct strategy, in which the C_1_ group is transferred to methylene-THF, a process that likely employs a corrinoid-dependent methyltransferase. The resulting methylene-THF is then channeled into the Wood-Ljungdahl pathway (WLP). Interestingly, Diel generates hydrogen during DCM mineralization to CO_2_ and chloride, which necessitates a syntrophic partnership with a hydrogen-consuming population (G. Chen, Kleindienst, et al., 2017). In contrast, ‘*Ca*. Formimonas warabiya’ ferments DCM via intracellular syntrophy to acetate and chloride (Holland et al., 2021; Wiechmann et al., 2020) and axenic Defo cultures ferment DCM to acetate, formate and chloride (G. Chen, Murdoch, et al., 2017).

Due to the genetic intractability of anaerobic DCM-degrading bacteria, comparative genomic approaches were applied to unravel underlying conserved genes involved in DCM metabolism, which led to discovery of a novel conserved gene cluster. Proteomics applied to Diel and Defo grown with DCM supported a role of this gene cluster in anaerobic DCM metabolism. Tracing this gene cluster by analyzing public metagenome datasets and performing targeted qPCR assays revealed the prevalence of this gene cassette and the potential DCM-metabolizing phenotype in various environmental systems. The findings suggest DCM formation and consumption in diverse natural ecosystems and provide new opportunities for assessing how global changes in climate and habitat patterns impact DCM emissions and associated ozone destruction.

## Methods

### Comparative genomics

Initial identification of homologous genes shared between the genomes of Diel and Defo was performed by BLASTP-based reciprocal best hit (RBH) analysis within the Integrated Microbial Genomes (IMG) system (I.-M. A. Chen et al., 2019) and by using GView (Petkau et al., 2010). The two homologous gene clusters present in the Diel genome were manually delineated. A homologous gene cluster was identified in the genome of ‘*Ca*. Formimonas warabiya’ by application of local BLASTP searches (Altschul et al., 1990). Functional annotations (COG, pfam, KEGG, TIGRFAM) for Defo and Diel genes were obtained from the IMG system, while those of ‘*Ca*. Formimonas warabiya’ were assigned using the WebMGA server (Wu et al., 2011) for COG (Tatusov et al., 2000), pfam (El-Gebali et al., 2019), and TIGRFAM (Haft et al., 2012) annotations and GhostKOALA (Kanehisa et al., 2016) for KEGG annotations (Kanehisa et al., 2017).

### Metagenome searches

All 18,314 metagenomes in the IMG database publicly available as of January 7, 2020 were subjected to BLASTP query with the Defo MecE protein sequence with a minimum bit score cutoff of 150 (approximately 40% identity). The resulting protein set was further filtered by applying a RBH criterion, retaining only proteins whose closest BLAST-P hit to the IMG genomes database was found among the MecE sequences located in the putative *mec* gene cassettes. The candidate metagenome MecE homologs were then further filtered by retaining only proteins encoded by genes co-localized with at least one other *mec* cassette protein, applying the same RBH criterion. All proteins encoded by genes located on scaffolds where the *mec* protein homologs were identified were downloaded, subjected to local BLASTP query using the ten *mec* gene cassette proteins from Defo, and plotted using GenoPlotR and custom R scripts (Guy et al., 2010). Gene copy per genome for metagenome *mec* cassettes, provided in Dataset S1, were calculated by dividing the read depth of the corresponding scaffold by an average read depth of ten single copy conserved protein-encoding genes (ribosomal proteins L11 (COG0080), L1 (COG0081), L3 (COG0087), L4 (COG0088), L2 (COG0090), L22 (COG0091), L5 (COG0094), L15 (COG0200), L10 (COG0244), and L29 (COG0255)).

### Phylogenetic reconstruction

The top 20 most similar proteins to each of the proteins encoded by each of the Defo genes located on the *mec* cassette were obtained by searching the IMG genome database using BLASTP with a confidence threshold of 1e-5, except in the case of MecC, for which the threshold was 1e-2. All *mec* cassette genes located in both genomes and metagenomes were aligned and subjected to phylogenetic reconstruction alongside the top 20 most similar genes located in microbial genomes in the IMG database. Proteins encoded by genes from metagenomes were clustered at 80% similarity using CD-hit (Li & Godzik, 2006). Sequences were aligned using MAFFT G-INS-I with 1,000 maximum iterations (Katoh & Standley, 2013), trimmed using trimAl-gappyout (Capella-Gutiérrez et al., 2009) and subjected to phylogenetic reconstruction using FastTree2 maximum-likelihood estimation (Gamma-LG model) (Price et al., 2010). The resulting Newick tree files were visualized using the Interactive Tree of Life (Letunic & Bork, 2016).

### Preparation of Diel and Defo cultures for global proteomics

The DCM-degrading consortium RM harboring ‘*Ca*. Dichloromethanomonas elyunquensis’ (Diel) and the axenic culture *Dehalobacterium formicoaceticum* (Defo) were grown in triplicate in 100 mL of anoxic mineral basal salt medium with 0.2 mM sodium sulfide, 0.2 mM L-cysteine (Löffler et al., 2005) and 30 mM bicarbonate (pH 7.3) under a headspace of N_2_/CO_2_ (80:20, vol/vol) with 156 µmol (10 µL) of DCM as the sole energy source. Cultures were initiated with a 5% (vol/vol) inoculum, incubated at 30°C in the dark without agitation, and provided one additional feeding of DCM once the initial amendment was consumed. Biomass for (meta)proteomic analyses was collected after 2 weeks of incubation when approximately 95% of the second DCM feedings were consumed. Culture suspensions were passed through Sterivex™ 0.22 µm membrane filter units (EMD Millipore Corporation, Billerica, MA, US) to capture cells. The outlet of the filters were capped, and 1.5 mL of boiling SDS lysis buffer (4% SDS in 100 mM Tris/HCl buffer, pH 8.0) were added to each of them. Filter unit inlets were then capped and placed in a laboratory rocker for 1-hour at room temperature. The SDS lysis buffer was removed by connecting 3 mL plastic syringes to the inlets of the cartridges, and then holding the syringes and filter units vertically and pushing air into each cartridge in order to withdraw as much lysate as possible by back pressure. In addition, filters were rinsed once more with 0.5 mL of fresh SDS lysis buffer. Lysate mixtures were centrifuged at 21,000 g for 15 mins and the clean protein supernatant transferred to fresh Eppendorf plastic tubes. Proteins were precipitated with trichloroacetic acid (TCA), denaturated in 8 M urea, reduced with dithiothreitol (DTT), alkylated with iodoacetamide (IAM), and digested with sequencing grade trypsin (Promega, 1:50 trypsin-to-protein [wt/wt]) (Yang et al., 2012). Protein concentrations were estimated with the BCA assay (Pierce Biotechnology, Waltham, MA, US) and crude protein and peptide extracts were stored at -80°C for subsequent LC-MS/MS analysis.

### Global proteomics analyses of Diel and Defo cultures

Global proteomics analyses were performed with an Orbitrap Q Exactive Plus mass spectrometer (Thermo Fisher Scientific, Waltham, MA, US) equipped with a nano-electrospray source (ESI) interfaced with a Proxeon EASY-nLC™ 1200 system. Peptides (2 µg) from each sample were suspended in solvent A (2% acetonitrile / 0.1% formic acid) and injected onto a C18 resin 75 μm microcapillary column (1.7 μm, 100Å, Phenomenex). Separation was accomplished at a constant flow rate of 250 nL/min with a 90-minute gradient from 2 to 30% solvent B (0.1% formic acid / 80% acetonitrile) followed by an increase to 40% solvent B within 10 minutes. Tandem mass spectrometry data (MS/MS) were collected using the Thermo Xcalibur software version 4.2.47 with similar parameters as reported before (Ganusova et al., 2021). Raw spectral files were searched against protein databases from the IMG annotated genomes of enrichment culture RM (which contains Diel) and Defo (IMG genome IDs 3300005804 and 2811995020, respectively), to which common laboratory contaminant proteins were appended. For standard database searching, the peptide MS/MS data was searched using Proteome Discoverer v2.4. The MS/MS data were searched using the SEQUEST HT algorithm (Eng et al., 1994) which was configured to derive fully tryptic peptides with the following settings: Maxium of 2 missed cleavage sites per peptide, minimum peptide length of 2, MS1 mass tolerance of 10 ppm and a MS2 tolerance of 0.02 Da. In addition, carbamidomethylations on cysteines (+57.0214 Da) and methionine oxidations (+5.9949 Da) were searched on peptides as static and dynamic modifications, respectively. Peptide spectrum match (PSM) confidence was evaluated with Percolator (Käll et al., 2007). PSMs and peptides were considered identified at a *q value* of < 0.01. Abundance values were converted to log2 values for ease of visualization. The IMG gene IDs of detected culture RM proteins were mapped to proteins contained in the IMG annotated genome of Diel (IMG genome ID 2627853586, Dataset S2).

### DCM measurements

DCM was quantified by manual headspace injections (0.1 mL) into an Agilent 7890 gas chromatograph (GC) (Santa Clara, CA, USA) equipped with a DB-624 column (60 m length, 0.32 mm i.d., 1.8 mm film thickness) and a flame ionization detector (FID). To analyze DCM concentrations in groundwater, 1-mL samples were collected, immediately transferred to sealed 20-mL glass vials, and the DCM concentration determined in the headspace. Aquoeus phase concentrations were determined using a dimensionless Henry’s law constant of 0.0895 (Gossett, 1987).

### Environmental samples

Anaerobic digester sludge was collected from two wastewater treatment plants, one located in Knoxville (KUB) and the other in Lenoir City (LC), TN. Groundwater samples from six monitoring wells representing within plume, fringe and outside locations at a DCM-contaminated site were obtained from CDM Smith (Wright et al., 2017). The groundwater samples were shipped with an overnight carrier in a cooler with ice and analyzed immediately upon receipt. Frank Stewart (Montana State University) provided archived DNA samples from two vertical transcects from the ETNP OMZ.

### DCM enrichments

Microcosms were established in 160-mL glass serum bottles containing 98 mL of anoxic mineral basal salt medium amended with 156 µmol (10 µL) of DCM. The microcosms were seeded with 2 mL of digester sludge, and additional DCM feedings occurred upon the depletion of DCM. Microcosms showing DCM degradation were sequentially transferred to fresh anoxic medium with DCM as the sole electron donor with an inoculation volume of 3 mL. After eight consective transfers, solids-free enrichment cultures were obtained that degraded DCM under anoxic conditions. DNA samples were extracted from the new DCM enrichment cultures and used to examine the presence of *mecE* and *mecF* genes by qPCR.

### DNA extraction

DNA extraction from 1 mL anaerobic digester sludge was performed using the DNeasy PowerSoil DNA extraction kit (Qiagen, Valencia, CA). DNA from Defo and Diel cultures was extracted from 5 mL culture suspensions collected onto 0.22 µm Durapore membrane filters (Millipore, Cork, Ireland) and DNA using the DNeasy PowerLyzer PowerSoil DNA extraction kit (Qiagen, Valencia, CA) following the manufacturer’s instructions.

Biomass from groundwater samples (950 mL) was collected on Supor® 0.2 µm membrane filters (Pall Lab., Ann Arbor, MI). Each filter was cut in half using a sterile scalpel and each piece was placed into a separate bead-beating tube for extraction with the DNeasy PowerLyzer PowerSoil DNA kit. The extracted DNA was concentrated with the Zymo DNA Clean and Concentrator-25 Kit (Zymo Research, Irvine, CA). DNA concentrations were determined with fluorometry and DNA was stored at -80°C until qPCR analysis.

### Primer design, PCR and qPCR analyses

Primer sets were developed for both *mecE* and *mecF* genes. The design was based on the target gene alleles identified in the genomes of Diel, Defo, Dcmf, and strain UNSWDHB and the most similar homologs from peat bog metagenomes. Additional primers were designed based on the most common *mecE* and *mecF* alleles identified in Eastern Pacific OMZ metagenomes (IMG genome Ga0066828, gene IDs 100177932 [*mecE*] and 100027434 [*mecF*]). The respective target gene sequences were aligned using ClustalW and primer sets were designed using the Primer 3 plug-in in Geneious R11.0.2 (Kearse et al., 2012) for PCR and SYBR qPCR assays. The primer sequences were blasted against NCBI nr database using the Primer BLAST program to verify specificity of the assays. The primers were obtained from a commercial supplier (Integrated DNA Technologies, Coralville, IA). For quantification of total bacterial 16S rRNA genes, previously reported Bac1055YF/Bac1392R (Ritalahti et al., 2006) and EUB338F/EUB518R primers were used (Lane, 1991; Muyzer et al., 1993). Primer sequences are listed in Table 1.

**Table 1.**
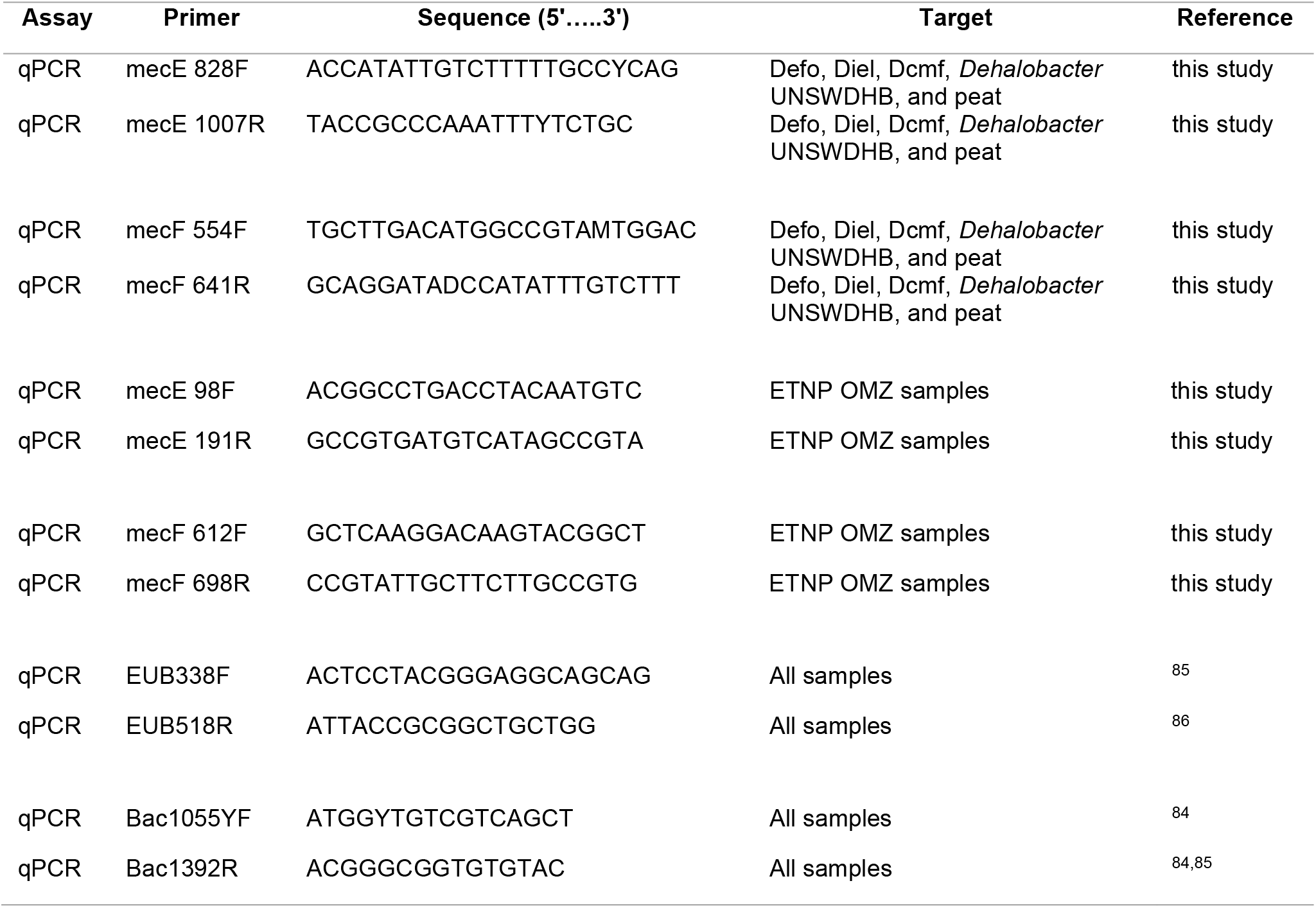
Novel primers used for qPCR. The column “Target” refers to the source of the template alleles used to design and validate the primers. peat; *mec* gene homologs derived from peat metagenomes.

qPCR was performed in 10-µL volumes consisting of 5 µL 2X Power SYBR Green PCR Master Mix (Applied Biosystems, Foster City, CA), 0.5 µL of each primer (final concentration of 300 nM) and 2 µL of template DNA (undiluted, 1:10 and 1:100 dilutions). qPCR analysis was conducted using a QuantStudio 12K Flex Real Time qPCR System (Life Technologies, Carlsbad, CA) and the thermocycling program was followed as initial step of 10 min at 95°C, 40 cycles of 15 sec at 95°C and 1 min at 60°C. Specific amplification was confirmed by melt curve analysis and agarose gel electrophoresis as described (Hatt & Löffler, 2012). Standard curves were established with linear GeneArt DNA fragments of target genes. Standards were run as triplicate on each plate using ten-fold dilution series in the range of 10^1^ to 10^8^ gene copies/µL. The amplification efficiencies, linear dynamic range, slope, Y-intercept and R^2^ values are listed in Dataset S3. Amplification efficiencies (AE) were calculated using the equation 10^(−1/slope)^ – 1.

### Metatranscriptomics

Unprocessed Illumina metatranscriptome sequencing data generated from groundwater from a DCM contamination plume was provided by CDM Smith. This same site was previously the subject of 16S rRNA gene amplicon library analysis (Wright et al., 2017). Raw data were trimmed using Trimmomatic v0.35 with a 6:25 sliding window quality trim, Illumina adapter read-through contamination removal, and final minimum length of 25 bp (Bolger et al., 2014). Trimmed reads were assembled *de novo* using Trinity v2.8.5 under default parameters (Grabherr et al., 2011). Prokaryotic ribosomal RNA genes were detected in the transcript contigs using barrnap v0.9-2 (Seeman, 2018) and removed using a custom R script employing Biostrings v2.5.4 (Pagès et al., 2019). Remaining transcripts were queried against a database consisting of all Defo *mec* genes using TBLASTX. Transcripts aligning to at least *mecE* with a bit-score of >1,000 (approximately 70% full-length amino acid alignment) were included. Transcript coverage was calculated using kallisto (Bray et al., 2016) and TPM values were calculated using the Trinity utility script align_and_estimate_abundance.pl.

## Results

### Anaerobic DCM degraders share a common gene cassette

Comparative analysis of the Defo and Diel genomes revealed that the eight most similar genes (and 10 of the top 25 most similar genes) in terms of percent predicted amino acid identity were located in genetic clusters. The genome of Defo harbors a single 10-gene cluster and Diel has two highly similar clusters A and B (Figure 2, Dataset S4).

**Figure 2.**
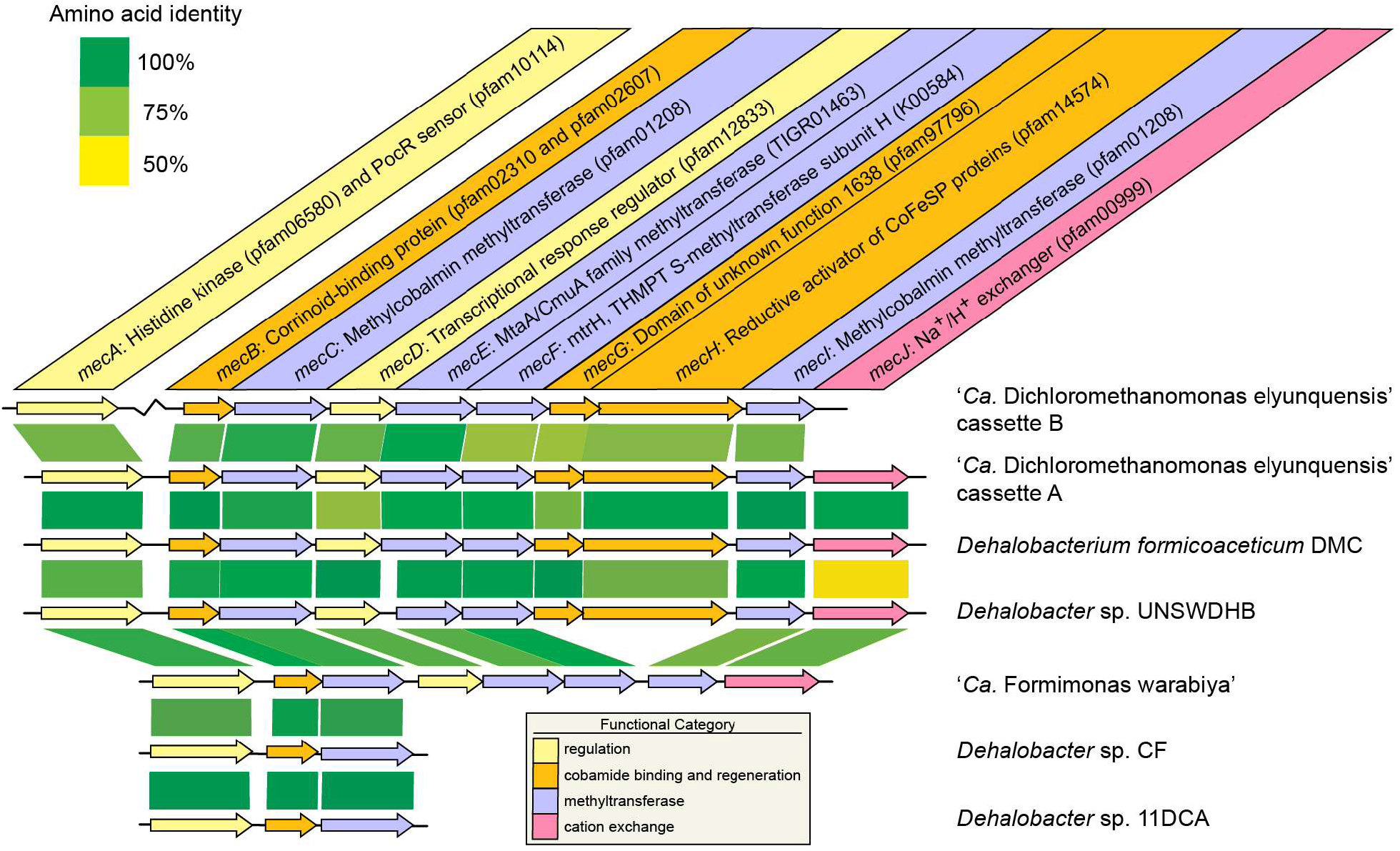
*mec* metabolic gene cassettes and close homologs identified in genomes. Shaded boxes represent BLASTP amino acid identity scores. The colors represent the general functional category of the encoded protein. Function was inferred from functional annotation systems, with priority given to the TIGRFAM and KEGG systems. GenoPlotR(*26*) was used to parse BLASTP results and gene coordinates to generate the figure.

The gene arrangement in these clusters is identical (i.e., syntenic) and predicted amino acid identities between clusters range from 79.6% to 99.7% (Figure 2). This novel, conserved 10-gene *mec* cassette harbors *mecA* thorough *mecJ* implicated in **<u>me</u>**thylene **<u>c</u>**hloride catabolism. A homologous gene cassette was also found in the newly described DCM degrader ‘*Ca*. Formimonas warabiya’ (Holland et al., 2019), although lacking *mecG* and *mecH*. Aside from *mecJ*, the only close homologs in any prokaryotic genome to any of the *mec*-encoded proteins are found in the genomes of three chloroform degrading bacteria; a homologous 10-gene cassette in *Dehalobacter* sp. strain UNSWDHB and partial gene cassettes in *Dehalobacter* sp. strain CF and *Dehalobacter* sp. strain 1,1-DCA (Figure 2). Functional annotation of the *mec* cassette gene products revealed a histidine kinase sensory protein and an associated regulatory protein (MecAD), an MtaA/CmuA methyltransferase (MecE), an MtrH methyltransferase (MecE), two methyltransferases of indeterminate function (MecCI), a corrinoid-binding protein (MecB), a cation transporter/antiporter (MecJ), a reductive activator of corrinoid proteins (MecH), and a protein with a conserved domain of unknown function (MecG) (Table 2 and Supplementary Text).

**Table 2.**
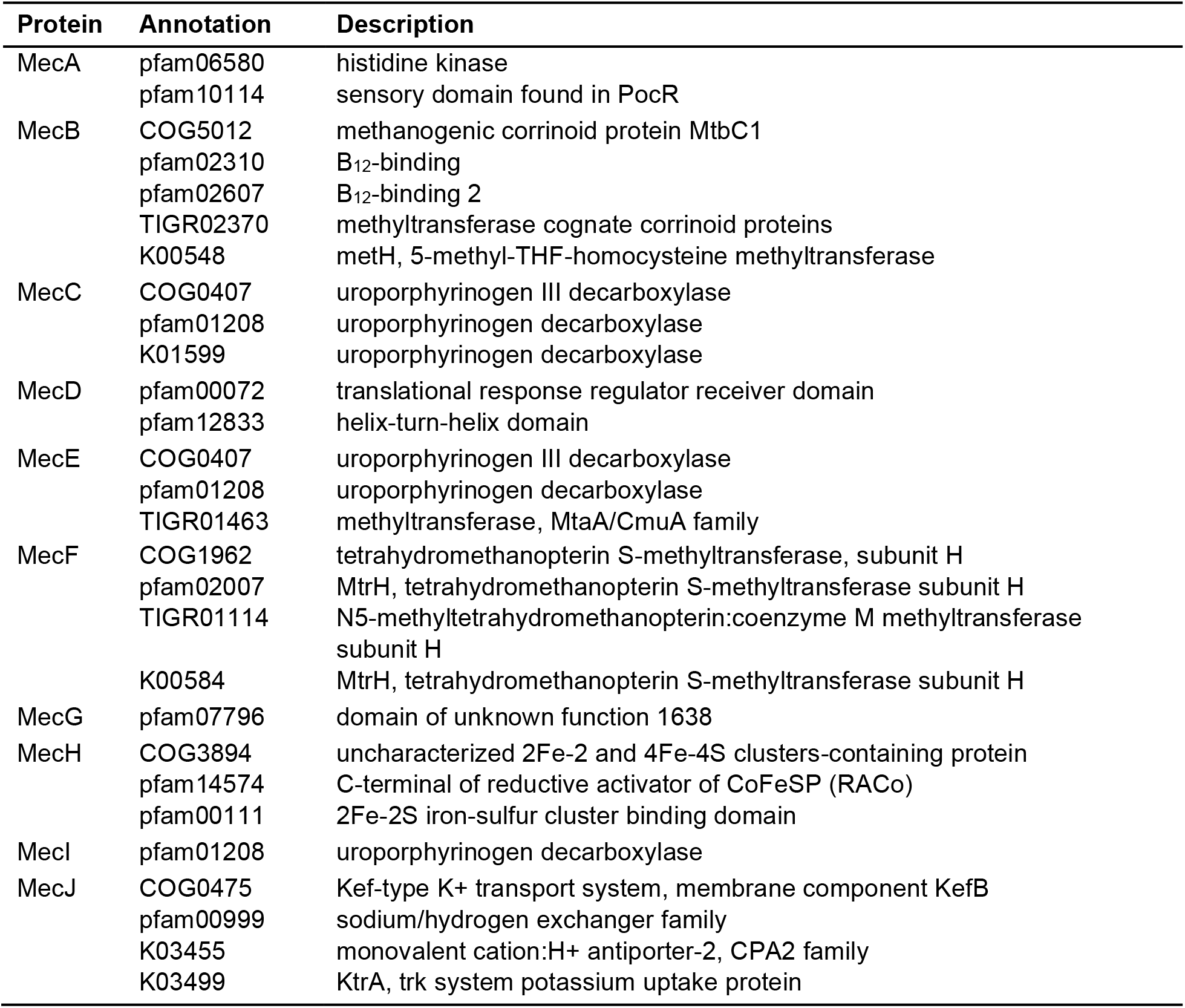
Consensus functional annotations of the putative *mec* cassette gene product proteins. COG, Clusters of Orthologous Genes; TIGRFAM, The Instutute for Genomic Research’s Database of Protein Families; pfam, Protein Familes; KEGG, Kyoto Encyclopedia of Genes and Genomes.

### Mec proteins are expressed during growth on DCM

When grown with DCM, a total of 1,781 proteins were detected in the axenic Defo culture (Dataset S5) and 1,743 proteins were detected in the metaproteome of mixed culture RM, 797 of which were assigned to Diel (Dataset S2). The majority of proteins encoded by the *mec* gene cassettes were detected in the proteomes of both DCM degraders (Figure 3, Table S1).

**Figure 3.**
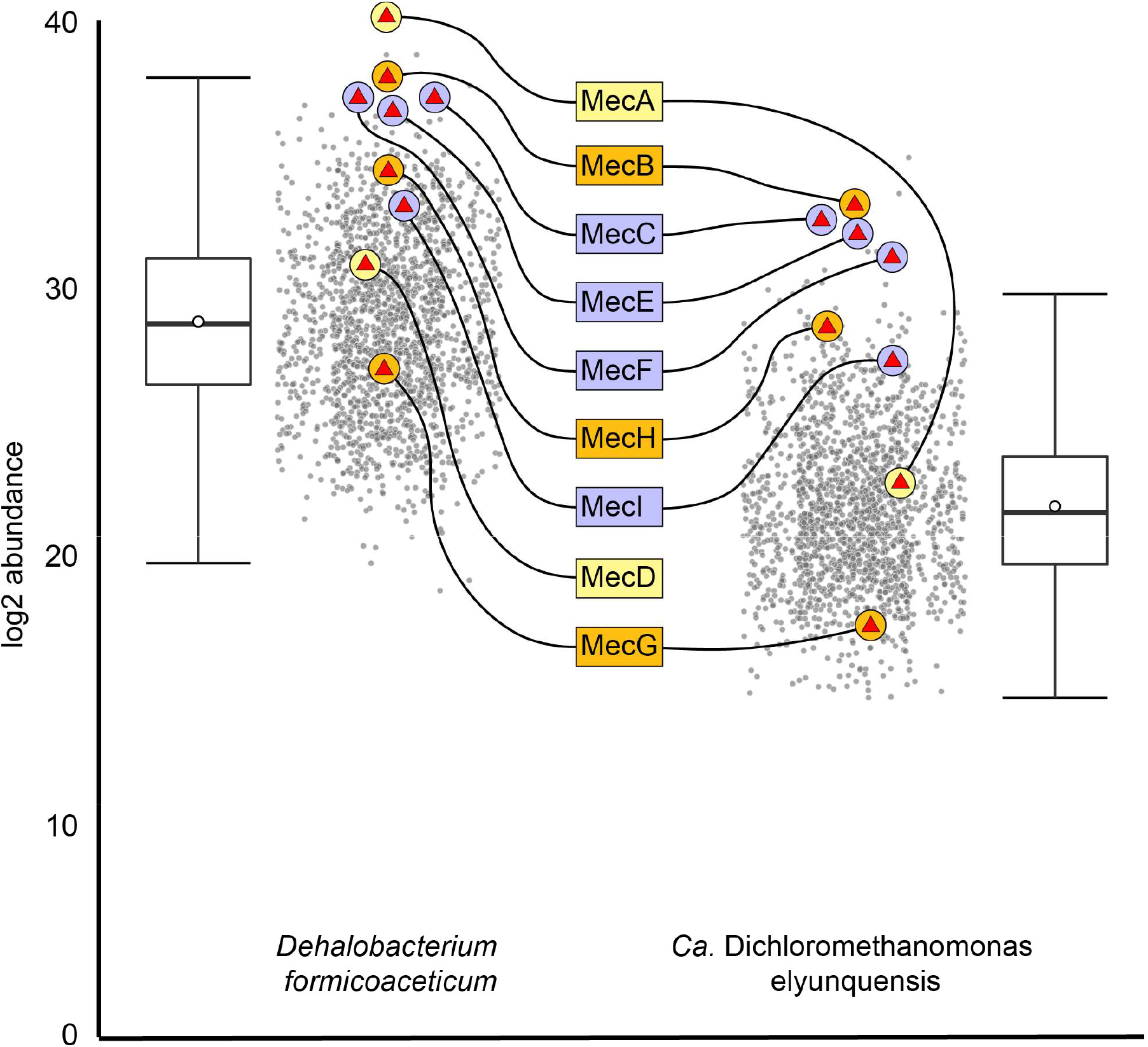
Mec protein abundance in the proteomes of Defo and Diel when grown anaerobically with DCM as the sole carbon and energy source. Box and whisker plots show median (central horizontal line), upper and lower quartile range (box), highest and lowest values excluding outliers (upper and lower whiskers) and mean value (open circle). Protein products of the *mec* cassette genes are labelled, with general functional category indicated by color (orange, corrinoid-related; purple, methyltransferase; yellow, regulatory).

In Diel, all proteins of *mec* cassette B were detected except for MecD and MecJ while in Defo, all but MecJ were detected. The corrinoid-binding protein MecB was the 2^nd^ and 3^rd^ most abundant protein in Defo and Diel proteomes, respectively. The three methyltransferases MecC, MecE, and MecF were in the top 1% most abundant proteins in both proteomes. The fourth methytransferase MecI and the corrinoid protein reductive activator MecH were all in the upper quartile of detected proteins. The sensor histidine kinase MecA was 1^st^ and 38^th^ most abundant in Defo and Diel proteomes, respectively. In neither case was MecJ detected, although its predicted eleven transmembrane alpha helices suggest strong association with the cytoplasmic membrane, likely hindering detection in the proteomics measurements (Vit & Petrak, 2017).

### DCM enriches for bacteria harboring mec genes

Targeted qPCR assays for the *mecE* and *mecF* genes did not yield quantifiable signals with template DNA extracted from anaerobic digestor sludge. Following enrichment with DCM using the same anaerobic sludge as inoculum, 1 × 10^6^ to 5 × 10^7^ gene copies/mL of both *mecE* and *mecF* were measured in transfer cultures, consistent with the observed consumption of DCM (Figure S1).

Groundwater samples from a DCM plume provided a unique opportunity to explore *mec* gene abundance and expression in response to varying DCM concentrations. *mecE*- and *mecF*-targeted qPCR assays yielded signals in a DCM dose-dependent manner, with ratios of *mec* gene copy number versus total bacterial 16S rRNA gene copy numbers ranging between 4% and 10% in groundwater from wells with 3.2 mg/L DCM or higher (Figure 4B).

**Figure 4.**
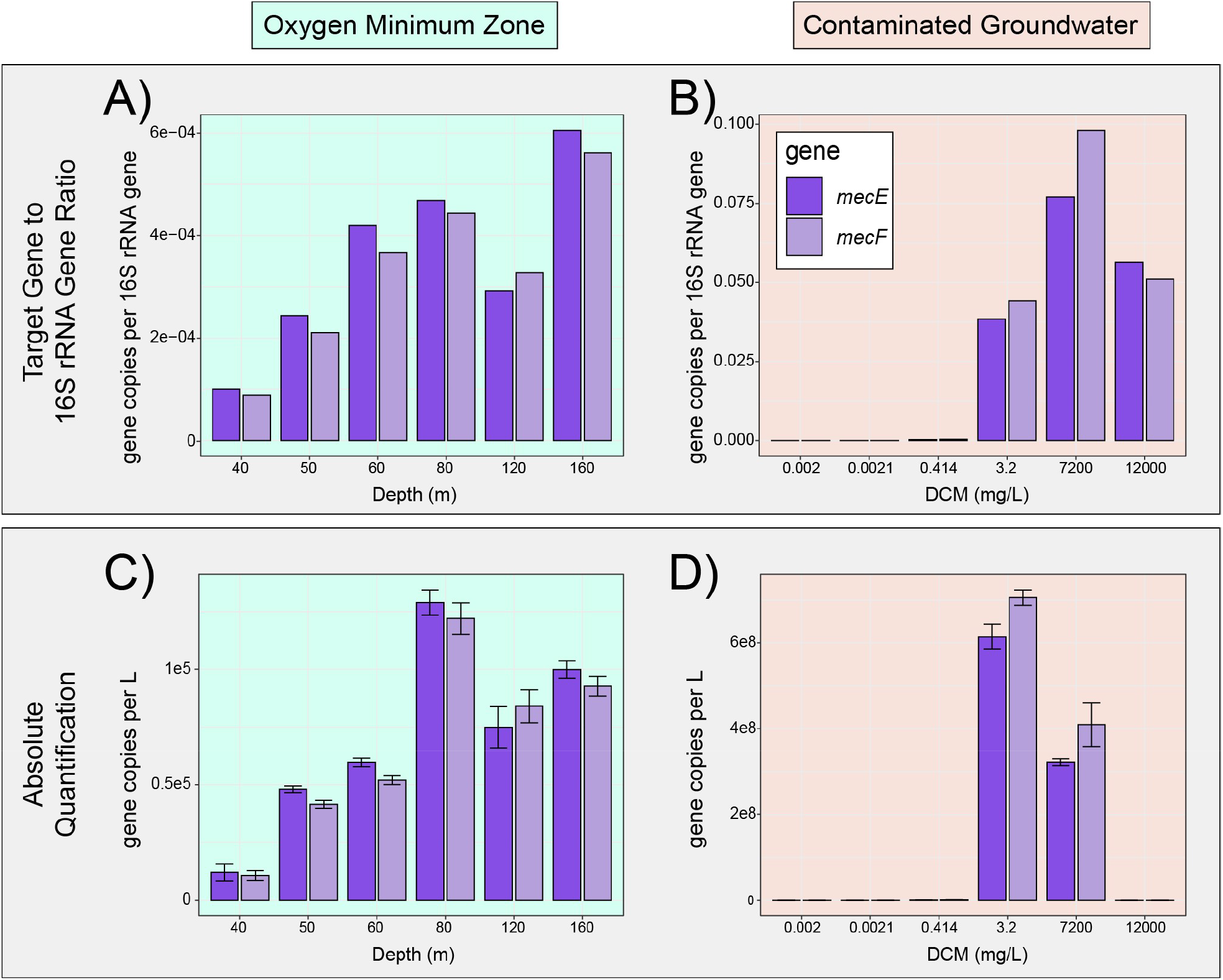
Quantitative PCR of *mecE* and *mecF* in the Eastern Pacific oxygen minimum zone (OMZ) and from groundwater samples from a DCM plume. A) and B) represent relative quantification results expressed as gene copies per 16S rRNA gene copy number for OMZ and DCM groundwater plume samples, respectively. Panels C) and D) represent absolute quantification (i.e., qPCR) results for OMZ and DCM groundwater plume

At lower DCM concentrations of 1.47 mg/L, the relative abundance of *mec* genes dropped to 0.029%, and in wells at the fringe of the plume with DCM in the low µg/L range, *mec* gene to 16S rRNA gene ratios dropped to 0.0004 – 0.0026% (Figure 4B). Absolute *mec* gene copy numbers followed this trend, except for samples collected from the well with the highest DCM concentration of 12 g/L, where lower abundances of bacterial and archaeal 16S rRNA genes were observed (i.e., 1.44 × 10^6^ versus 3.57 × 10^9^ − 1.60 × 10^10^ 16S rRNA genes per L at locations with lower DCM concentrations) (Figure 4D). Absolute target gene copy numbers and ratios of target genes to total bacterial 16S rRNA copy numbers both covaried with measured DCM concentrations, with no detections outside the plume and *mec* gene to 16S rRNA gene ratios of up to 10% within the plume, indicating that roughly 1 in 10 bacterial genomes in the plume harbored a *mec* cassette.

Analysis of metatranscriptomes, previously obtained for groundwater microbiomes collected from the same DCM-contaminated site, revealed expression of *mec* cassette genes. *mecE* and *mecF* were consistently identified on the same transcript, with relative expression reaching its highest value of 44.6 transcripts per million transcripts (TPM) in wells located in the plume fringes. *mec* cassette transcripts were also detected in groundwater samples in the core plume at relative expression levels up to 21.2 TPM (Table S2).

### Environmental distribution of the mec cassette

The search of over 18,000 metagenomes identified similar gene cassettes in 41 metagenomes from peatland, the deep subsurface, and marine systems (Figure 5, Figure S2). The SI provides gene and genome IDs and detailed BLAST-P results (Table S3, Dataset S6, Dataset S7).

**Figure 5.**
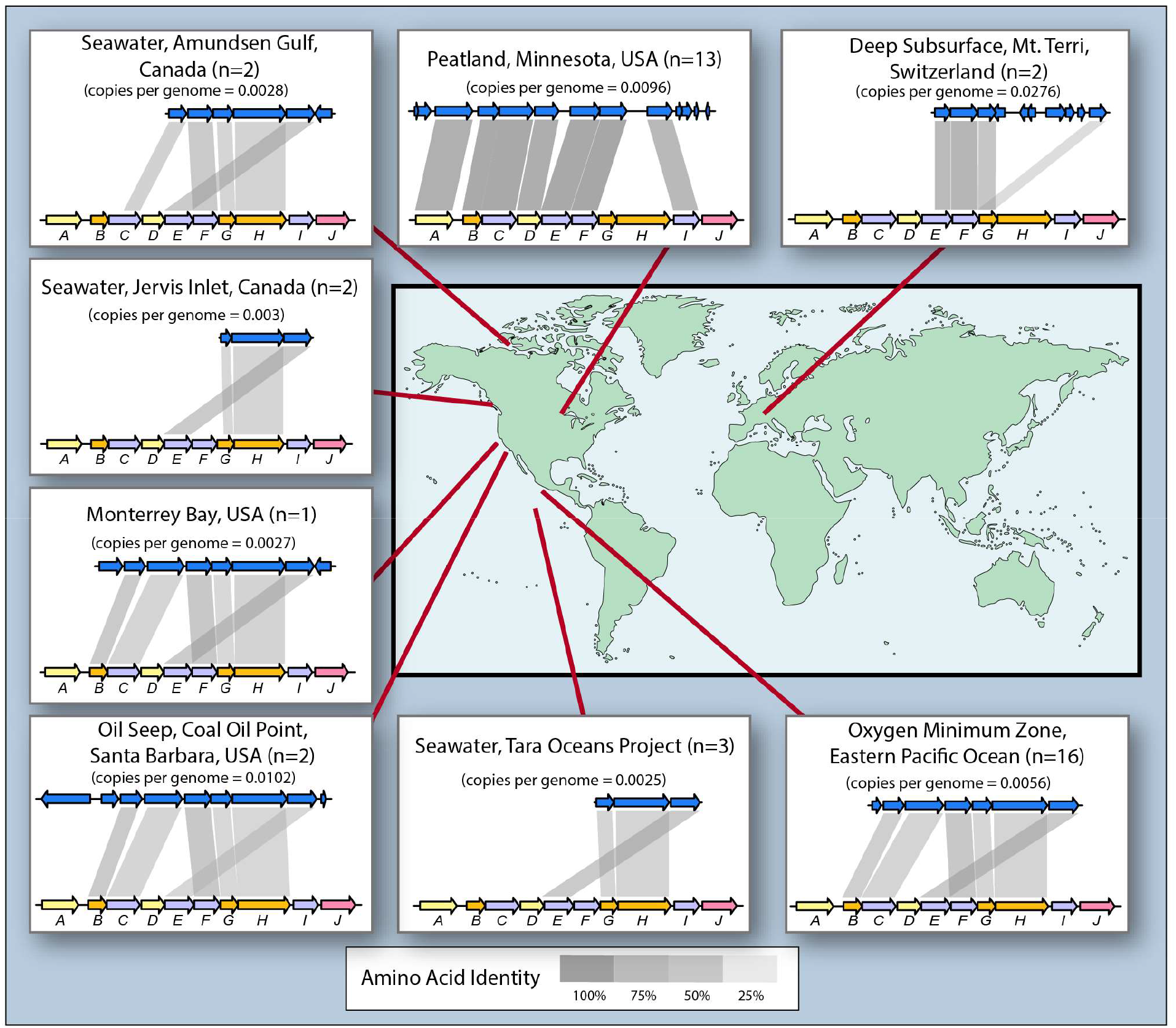
Distribution of 41 gene cassettes with homology and synteny to the Defo *mec* cassette. Identified in the IMG metagenome database, details provided in Dataset S7. Representative cassettes from the eight environments, in which the cassettes were identified, are shown. The red lines point to the approximate locations from where the metagenomes were derived. In each gene cassette illustration, the lower, multicolored gene cassette represents the *D. formicoaceticum mec* cassette, while the blue, upper illustration represents the gene cassette found in the indicated environmental metagenome. “n” indicates the number of individual samples, in which a syntenic cassette was identified, while “copies per genome” indicates the estimate of cassette copy per total genomes present in the metagenome, as determined by comparing cassette read depth to the average read depth of ten single-copy protein-encoding genes for the cassette shown. Shaded boxes show the BLAST-P-derived amino acid identity allowing comparisons of gene-product identities.

*mec* cassettes were identified in 13 of 117 metagenomes from ombrotrophic peat bogs in the Marcell experimental forest in Minnesota, USA. The *mec* cassettes were predominantly (i.e., 10 out of 13 gene cassettes) identified in metagenomes from samples collected between 1 – 1.5 m depth, where the pH was approximately 4.5 and anoxic conditions prevailed (Chris Schadt, personal communication). Two cassettes organized as *mecABCDEFI* displayed predicted amino acid identity scores to the Defo *mec* cassette genes above 80% (Figure 2) and close phylogenetic affiliation (Figure S3). This *mecABCDEFI* cassette was most prevalent, present in nearly 1% of all genome copies (i.e., approximately 1 out of 100 prokaryotic cells in the community harbor a *mec* cassette) (Dataset S7).

A total of 23 *mec* gene cassettes were identified in metagenomes from samples collected beneath the photic zone of the oceanic water column. The majority of cassettes was identified in metagenomes from the Eastern Pacific oxygen minimum zone (OMZ) at depths of 150 m to 400 m, wherein oxygen was below the limit of detection (Thamdrup et al., 2019). Additional cassettes detected in metagenomes derived from three coastal sites, Monterrey Bay, CA, Jervis Inlet in British Columbia, Canada, and Amundsen Gulf, in Arctic northern Canada, and from an open ocean sample from the Eastern Tropical North Pacific (ETNP). The marine *mec* cassettes were syntenic, with the order *mecBCFGHE* (Figure 5). *mecE* and *mecF* were the most similar to the corresponding Defo homologs, with average amino acid identities of 49.7 - 51.6%. All of the marine *mec* genes were more closely related to one another and to the *mec* cassette genes of the characterized DCM degraders than to any other gene in assembled metagenomes or genomes available in NCBI or the IMG database. The marine *mec* gene clusters formed distinct, deeply branching clades (Figure S3). The highest marine *mec* cassette occurrence was observed in a sample from Coal Oil Point, CA, a natural marine petroleum seep area (present in 1.02% of total genome copies) (Dataset S7). qPCR applied to water column samples from two ETNP OMZ locations detected *mecE* and *mecF* at a higher frequency (9 out of 11) than they were found in the ETNP OMZ metagenome assemblies (16 out of 90), with target gene-to-bacterial 16S rRNA gene ratios of 0.01 – 0.06% (Figure 4, Dataset S8).

Evidence was obtained for the presence of *mecEFG* in anoxic porewater from a hydrogen-amended borehole in Opalinus Clay rock situated 300 m beneath Mt. Terri, Switzerland (Bagnoud et al., 2016). The amino acid identities for the putative methyltransferases MecE and MecF were 64.0 and 66.7%, respectively. Accordingly, phylogenetic analysis revealed close relationships with *mecE* and *mecF* of the known DCM degraders (Figure S3). The deep subsurface *mec* cassettes were present in 2.8% of total estimated genome copies.

### Genomes of chloroform-respiring organisms harbor mec gene orthologs

The *mec* gene orthologs comprise cohesive, deeply branching clades (Figure S3). Aside from *mecJ*, no close homologs (i.e., proteins with >35% amino acid identity) to the *mec* genes are found in any publicly available bacterial or archaeal genomes, with three notable exceptions. Homologs of *mecA, mecB*, and *mecC* are found on genomes of the chloroform (CF) respirers *Dehalobacter* sp. strain CF and *Dehalobacter* sp. strain 11DCA, both of which were reported to generate DCM as an end product during growth with CF as electron acceptor (Grostern et al., 2010). In both of these genomes, the *mecABC* homologs are found immediately adjacent to the CF reductive dehalogenase and anchor protein encoding genes *cfrAB*, suggesting functional association between the two gene clusters. In the third case, a complete 10-gene *mec* cassette is located on the genome of *Dehalobacter* sp. strain UNSWDHB (Figure 2), also a CF-respiring organism that lacks the ability to utilize DCM (Wong et al., 2016). Close inspection of the *mec* cassette of strain UNSWDHB reveals that *mecE*, implicated in the initial chloromethyltransfer reaction, is truncated at the 5’ end, which is projected to lead to a ∼70 amino acid shorter protein (Figure S4), consistent with a loss of function (Wong et al., 2016).

## Discussion

### Identification DCM biomarker genes

The 10 gene *mec* cassette was initially identified by comparative genomic analyses between the anaerobic DCM degrading bacteria *Dehalobacterium formicoaceticum* (Defo) and ‘*Ca*.’ Dichloromethanomonas elyunquensis (Diel). A third highly similar and syntenic gene cassette was also identified in the newer DCM degrader ‘*Ca*.’ Formamonas warabiya (Dcmf). This high degree of gene identity and gene order among DCM degraders and the near total absence of any closely-related gene cassette in any other bacterial genome placed strong suspicion on this gene cassette as being involved in anaerobic DCM catabolism. To confirm the association of these genes with anaerobic growth on DCM, additional enrichments were generated from municipal wastewater in Eastern Tennessee, which led to an increase in the abundance of *mec* genes from undetectable to levels similar to those seen in Defo and Diel cultures (1e6 to 1e8 gene copies per mL). Mec proteins were also among the most abundant proteins in the proteomes of DCM-grown Defo and Diel. Furthermore, it was observed that *mec* genes are enriched in the bacterial community at a DCM-contaminated groundwater site and were expressed in proportion to DCM concentration. The multiple lines of evidence espouse a strong and exclusive coordination between anaerobic DCM metabolism and *mec* cassette gene, transcript and protein abundance.

### Functional annotations of the mec cassette are consistent with a DCM dehalogenating methyltransferase system

The DCM-degrading bacteria are strict anaerobes and no genes consistent with aerobic DCM catabolism by a GST (e.g., *dcmA*) were identified in the genomes of strains Defo, Diel, or Dcmf. Previous biochemical studies on Defo demonstrated that DCM is channeled into the WLP via methylene-THF catalyzed by a corrinoid methyltransferase (G. Chen et al., 2018, 2020; A Mägli et al., 1998; Andreas Mägli et al., 1996). Guided by functional annotation (Table 2) and comparison to analogous metabolic systems, the proteins encoded by the *mec* cassette genes may comprise a DCM catabolic pathway that is compatible with available biochemical evidence (Figure 6).

**Figure 6.**
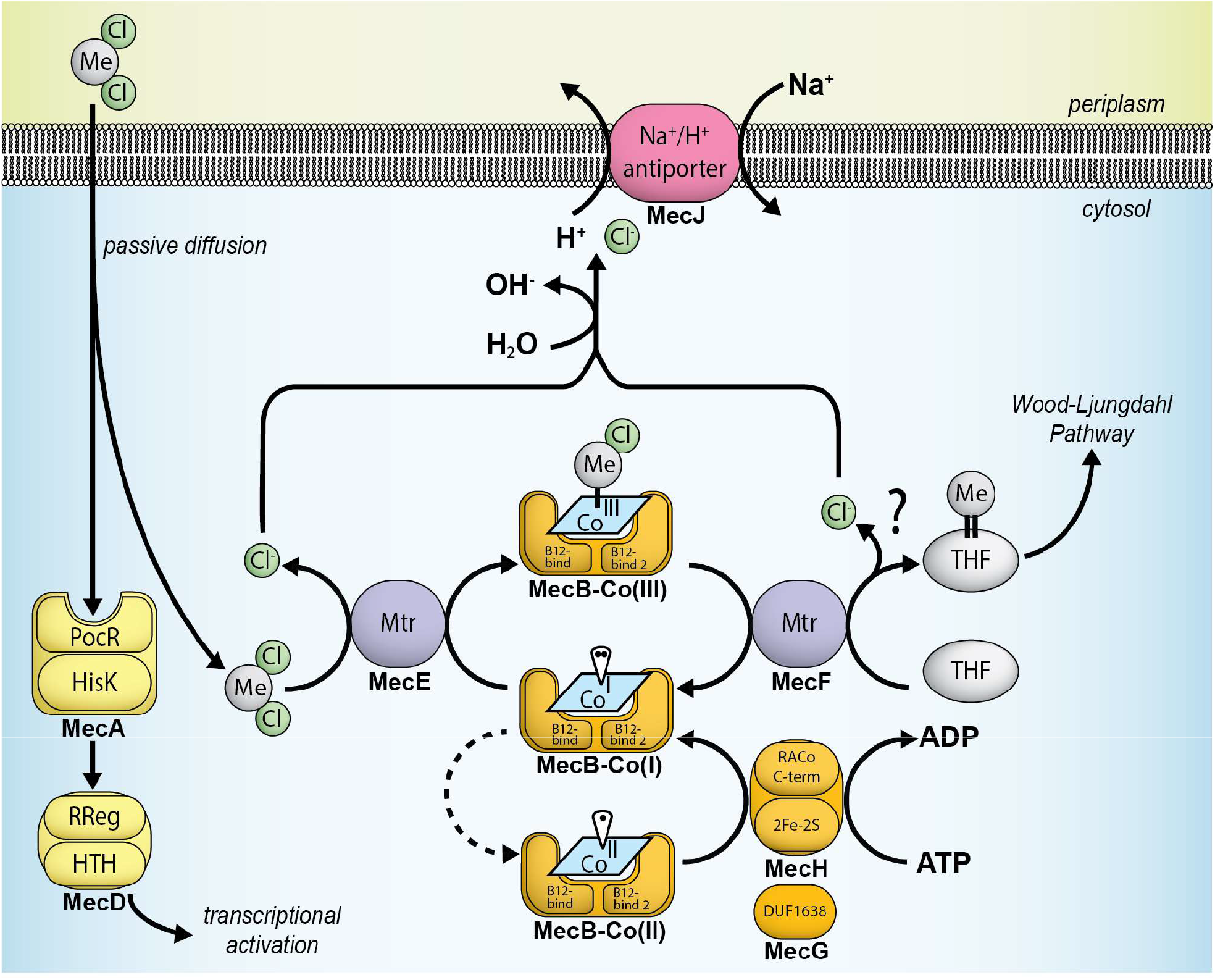
Proposed pathway for anaerobic DCM catabolism catalyzed by proteins encoded by the *mec* cassette identified in this study. Proteins with regulatory (yellow), methyltransferase (purple), B_12_-related (orange), transport (pink) functions are depicted. PocR, sensory domain found in PocR; HisK, histidine kinase; RReg, translational response regulator receiver domain; HTH, helix-turn-helix domain; Mtr, methyltransferase; B12-bind, B_12_-binding domain; RACo, reductive activator of CoFeSP; THF, tetrahydrofolate. Mechanistic understanding about the release of the second chlorine substituent has not been obtained, as indicated by the question mark. Experimental CSIA data suggest that different mechanisms operate in different strains (*40*).

The core system involved in C_1_ group transfer is likely composed of two methyltransferases, MecE and MecF, and a corrinoid-binding protein, MecB. MecE is a member of the MtaA/CmuA family (TIGR01463), an ortholog family defined by methylamine, methanol and chloromethane methyltransferases. The relationship with such methyltransferases, especially the chloromethane dehalogenase CmuA of the aerobic bacterium *Hyphomicrobium* sp. strain MC1 (Borodina et al., 2004), renders MecE a candidate for catalyzing the initial chloromethyltransfer reaction to remove one chlorine substituent from DCM. Chloromethane is dechlorinated by a corrinoid-dependent methyltransferase (CmuA), suggesting MecE could perform an analogous single dechlorination of DCM during C_1_ group transfer to the corrinoid-binding protein MecB. Such a reaction would yield a hypothetical chloromethyl corrinoid protein (Figure 6). Methyltransferase MecF is a member of the MtrH enzyme family responsible for transfer of a corrinoid-bound methyl group to the organic methyl carrier tetrahydromethanopterin (THMPT), a cofactor for C_1_ transfer analogous to tetrahydrofolate (THF). MecF is the best candidate for the final methyltransfer reaction to THF generating methylene-THF, for which biochemical evidence exists (A Mägli et al., 1998). This methyltransfer reaction could lead to C-Cl bond cleavage and release of the second chlorine substituent. Alternatively, another methyltransferase, such as MecC or MecI, or another enzyme not necessarily conserved among all anaerobic DCM degraders, is involved in the release of the chlorine substitutent. Previously, C and Cl stable isotope measurements provided evidence for distinct dechlorination mechanisms in anaerobic DCM degraders, and ^13^C-tracer experiments demonstrated different end products, even though both Diel and Defo employ the WLP for DCM catabolism (G. Chen et al., 2018, 2020).

Features of the remaining *mec* cassette gene products suggest roles in support of the dehalogenating C_1_ group transfer system and gene expression regulation. MecGH share domain organization with proteins implicated in activating the corrinoid-binding protein involved in methyltransferase-mediated methionine synthesis in a variety of bacteria (Price et al., 2018). Thus, MecGH may function as a reductive activator for MecB. MecJ is predicted to be a monovalent cation:proton antiporter, possibly supporting DCM catabolism by maintaining acid-base homeostasis (E Pinner, 1994; Roosild et al., 2010), counteracting acidification of the cytosol by hydrochloric acid (i.e., proton) generation during dechlorination of DCM (Ferguson et al., 2000). Co-localization of dehalogenase genes (e.g., reductive dehalogenases, haloacid dehalogenases) and *mecJ* homologs is common among related organisms (Table S4), and the gene cluster *cmuABC* for aerobic chloromethane utilization is 6 kb upstream and in the same orientation of a gene (IMG ID 650984269) encoding a putative chloride carrier/channel (ClC) protein, a protein family also implicated in acid-base homeostasis (Iyer et al., 2002). The histidine kinase MecA likely represents a DCM sensor kinase that regulates the DNA-binding protein MecD following detection of DCM via the PocR domain (Anantharaman & Aravind, 2005). The high abundance of MecA sensor kinase in Defo and Diel proteomes (1^st^ and 38^th^ most abundant protein, respectively) may suggest an additional or alternative role. In the marine *mec* cassettes, the regulatory genes *mecA* and *mecD* were not detected; however, in two cases, homologs of *dcmR*, which encodes the DCM sensor and regulatory protein utilized by aerobic DCM degraders (La Roche & Leisinger, 1991), were identified directly adjacent to the *mec* cassette (Figure 7) and could fulfill regulatory functions.

**Figure 7.**
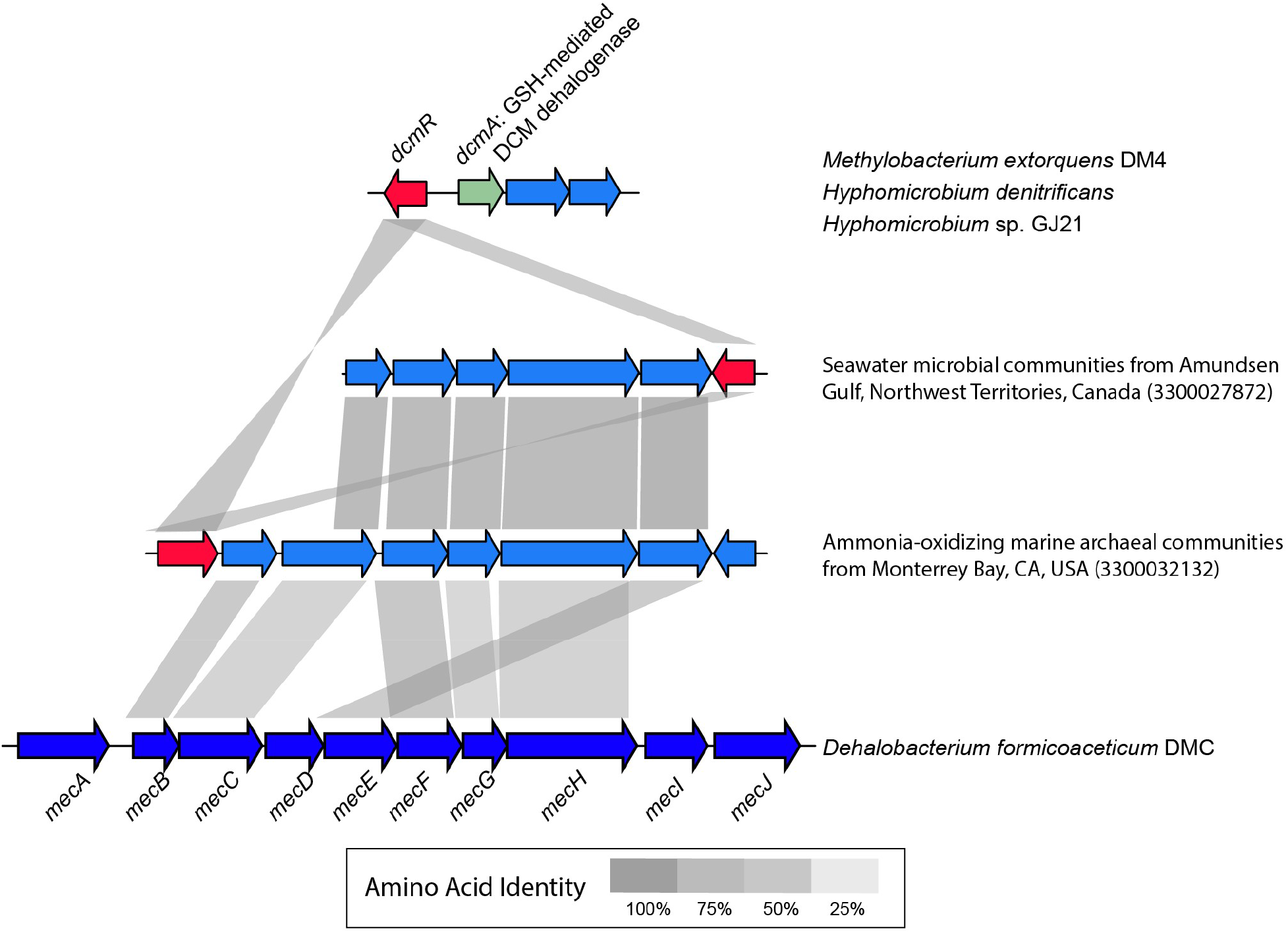
Two mec cassettes identified in metagenomes (from Monterrey Bay and Amundsen Gulf, CA) are flanked by genes, colored red, encoding proteins annotated as DcmR sensory domain and regulator of DCM dehalogenase DcmA (KEGG K17071). The products of these genes share 70.6% amino acid identity and are 44.6% - 46.4% identical to DcmR found in *Methylbacterium extorquens* strain DM4. The indicated gene cassette (*dcmRA*) encodes glutathione-mediated DCM dehalogenation.

The proposed involvement of additional methyltransferases (i.e., MecC and MecI) is not unprecedented. For example, the chloromethane utilization gene cluster in *Hyphomicrobium* sp. strain MC1 also contains a third putative methyltransferase encoding gene, *cmuC*, which is required for growth with chloromethane but whose function has yet to be revealed (Vannelli et al., 1999). Methanol-grown *Desulfotmaculum kuznetsovii*, which employs a similar methanol corrinoid methyltransferase system, also expresses three methyltransferases all located in the same gene cluster (Sousa et al., 2018). Additional details of the functional annotations of the *mec* cassette genes are provided in Supplementary Information.

Highly similar mec cassettes are broadly distributed in the environment. A search of public metagenomes led to identification of *mec* cassettes in disparate ecosystems, including peatland, marine oxygen minimum zones, and the deep subsurface. However, there is good reason to suspect each of these environments as being hotspots for DCM flux. Peat bogs have been demonstrated to be rich in chlorinated organic material (Biester et al., 2004) and have been identified as a source of halomethanes (Dimmer et al., 2001). The occurrence of halomethanes in subsurface rock formations has been demonstrated (Mulder et al., 2013), although information about quantities and fluxes is lacking. Marine systems are net producers of DCM and considered major emitters to the atmosphere (Gribble, 2010). In the water column, DCM concentrations peak along with chlorophyll concentrations (Ooki & Yokouchi, 2011), consistent with production by phytoplankton and the abiotic chlorination of planktonic iodo- and bromomethanes (Ooki & Yokouchi, 2011). DCM is detected beneath the photic zone (Ooki & Yokouchi, 2011), suggesting that mixing events carry DCM into deeper waters, or other sources of DCM exist in the deep ocean, possibly hydrothermal vents (Eklund et al., 1988; Isidorov et al., 1990; Jordan et al., 2000) or settling dead biomass (Wever & Barnett, 2017). Importantly, analysis of ETNP OMZ samples using targeted qPCR assays led to much high detection rate (9 of 11) than was obtained by metagenome analyses (16 of 90). This increase in frequency of detection is likely due to the higher sensitivity of qPCR compared to shotgun metagenomics (Suttner et al., 2020) and implies that the *mec* cassette is more broadly distributed in marine systems than the metagenomics survey results suggest.

### Implications of widespread anaerobic DCM degradation potential

DCM predates the anthropocene and the mechanisms underlying natural releases are far from fully characterized. Based on the available information, DCM is an energy source readily available to microorganisms in various environments, and the anaerobic microbial consumption of DCM is likely a major attenuation factor, eliminating DCM in anoxic systems prior to atmospheric release, and thus a relevant process for reducing DCM emissions. Environmental change, including global warming, has high potential to alter the flux of DCM with unpredictable consequences for the integrity of the ozone layer. Whether low oxygen marine systems and peat bogs, for example, are net producers or net sinks of DCM is currently unclear. OMZs are expanding at accelerating rates worldwide (Stramma et al., 2008), and uncertainty exists over the impact of environmental change on net DCM emissions. Likewise, climate change induced melting of permafrost in the northern hemisphere will create more active peat bogs, and warming of peat bogs is expected to mobilize recalcitrant carbon and stimulate the breakdown of accumulated organic materials (Gill et al., 2017), which will likely increase halomethane, including DCM formation. As is the case with low oxygen marine systems, the degree to which DCM production is counterbalanced by consumption in peat bogs is unknown. A widely distributed phenotype for anaerobic DCM catabolism is likely to affect DCM pools, fluxes and thus the global DCM budget, which until present considers atmospheric releases and abiotic stratospheric breakdown, but not microbial attenuation (i.e., sinks). The increased knowledge of microbial DCM catabolism offers opportunities to include relevant sink and attenuation terms and generate refined flux models with more predictive power.

## Acknowledgments

We thank F. Stewart and A. Bertagnolli for valuable discussions and providing Eastern Tropical North Pacific Oxygen Minimum Zone DNA samples for qPCR analysis. T. Macbeth, K. Sorenson, and R. Chenenko from CDM Smith graciously provided groundwater samples and metatranscriptomic data from a DCM-contaminated site. The authors acknowledge N. Ivanova and R. Seshadri from the Joint Genome Institute for providing invaluable insight into use of the IMG database.

## Data and Code Availability

All genomes, genes, and protein sequences studied are available in the IMG database (I.-M. A. Chen et al., 2019) under the specified ID numbers. Code used for data processing and figure creation is available via RPubs links located in the methods section. All spectral data collected from the Defo axenic cultures used in this study have been deposited in the MASSIVE and ProteomeXchange repositories with identifiers MSV000087235 and PXD025479, respectively. Spectral data collected from the Diel axenic cultures used in this study have been deposited in the MASSIVE and ProteomeXchange repositories with identifiers MSV000086520 and PXD022742, respectively. Scripts used to perform gene copy per genome copy calculations, pairwise gene cassette alignments, and to construct gene cassette synteny plots can be found at https://rpubs.com/rmurdoch/mec_cassette_abundance_and_synteny. Detailed description of gene phylogeny pipelines can be found at https://rpubs.com/rmurdoch/mec_cassette_trees. Full metatranscriptome analysis pipeline description and scripts used to operate programs can be found at https://rpubs.com/rmurdoch/mec_transcriptomes.

