## Supplemental Text, Figures, and Tables for "Global distribution of anaerobic dichloromethane degradation potential"

**This PDF file includes:**

Supplementary Text
Figs. S1 to S4
Tables S1 to S4
Legends for Datasets S1 to S8
Supplemental References

### Supplementary Text

Functional annotation of proteins encoded on the mec gene cassette

MecA and MecD. MecA contains two pfam domains, an N-terminal PocR (pfam10114) and a centrally located histidine kinase (pfam06580). A third domain was identified at the C-terminal, histidine kinase ATPase (HATPase, pfam02518), although the confidence of the assignment was low (8.12e-03). The pfam06580 and pfam02518 domains represent conserved regions of histidine kinase proteins which transmit signal to a response regulator while the PocR sensor domain contains a PAS-fold predicted to bind to small hydrocarbons such as 1,3-propanediol (Anantharaman & Aravind, 2005). MecD contains two conserved domains, pfam00072, a response regulator domain which receives signal from a sensor partner protein in two-component regulatory systems, and pfam12833, a DNA-binding helix-turn-helix domain which is found in many regulatory DNA binding proteins. Unlike typical sensor histidine kinases (Krell et al., 2010), MecA has no predicted transmembrane domains, nor does it contain any translocation signal peptides, implying cytoplasmic location.

MecB. MecB is the only protein encoded on the *mec* gene cassette with corrinoid binding domains (pfam02310 and pfam02607). MecB is annotated by both the COG and TIGR systems as being orthologous with corrinoid proteins found in *Methanosarcina* spp. Notably, the COG system places MecB specifically into the dimethylamine corrinoid methyltransferase protein MtbC1 group (COG5012). The corresponding TIGR02370 ortholog group is defined by trimethylamine, dimethylamine, and monomethylamine methyltransferases from *Methanosarcina* and a fourth protein, MtaF, for which methanol is the substrate.

MecE and MecF. The TIGR system places Defo MecE into the MtaA/CmuA subfamily of methyltransferases (TIGR01463). This group includes CmuA, a methyltransferase that is encoded by a chloromethane utilization gene cluster described in *Hyphomicrobium* species (McAnulla et al., 2001). MecF is consistently annotated as a tetrahydromethanopterin S-methyltransferase subunit H (COG1962, TIGR0114, pfam02007, K00584), the enzyme that transfers a methyl group to THF in canonical methanogenesis. MecF is a distant ortholog of CmuB from the same chloromethane utilization cluster (27.1% AA identity, 46.3% AA similarity); CmuB is responsible for transfer of the methyl group from methylcobalamin to a THF cofactor in chloromethane metabolism.

MecG and MecH. MecG is a member of the DUF1638 family (pfam07796). MecH contains two conserved domains, pfam14574 (C-terminal domain of RACo, formerly DUF4445) and a 2Fe2S iron-sulfur cluster assigned to pfam00111. These two genes commonly co-occur in genomes. DUF4445 genes have been found to be related to *ramA*, which encodes a protein that regenerates spent corrinoids in methanogens, utilizing ATP to reactivate (i.e. reduce) the cobalt cofactor that has entered the Co(II) to active Co(I) (Price et al., 2018). This pair of proteins is not found in the putative *mec* cluster in recently announced strain Dcmf.

MecC and MecI. Both of these proteins are annotated as non-specific methyltransferases (pfam01208, K01599, COG0407). MecC and Mecl sequences lack similarity to both to one-another to any characterized proteins.

MecJ. MecJ is designated by different functional annotation systems as a member of various families of cation transporters; Kef-type K+ transport system membrane component (COG0475 and K03499), Na+/H+ exchanger (pfam00999). The Transporter Classification Database (Saier et al., 2016) identifies MecJ as a monovalent cation:proton antiporter-2 (CPA2) family protein. MecJ is composed of 13 projected transmembrane helices (Figure S2); MecJ is the only protein within the cassette that contains any predicted transmembrane helices.

Estimation of global DCM release by wetlands

Measurements of DCM emissions from a coastal wetlands site ($2x10^{-8}$ $g/hr/m^2$ (Cox et al., 2004)) and recent estimates of total global wetlands area (Hu et al., 2017) leads to a an estimated global emission of 0.268 – 2.61 Gg/yr, while mangrove forests emit 1-2 Gg of DCM per year (Kolusu et al., 2018).

**Table S1.** Summary of *mec* cassette homolog proteomics data. "Gene.ID" indicates the IMG unique gene ID. "mec.protein" indicates the specific protein encoded on the *mec* cassette. "Rank" corresponds to the average relative abundance ranking within each organism's full proteome results. "log2.mean.abund" is the $\log_2$ transformed mean of the calculated protein abundance.

| *Dehalobacterium formicoaceticum* | | | |
|---|---|---|---|
| **gene.ID** | **mec.protein** | **rank** | **log2.mean.abund** |
| 2812836286 | MecA | 1 | 40.66 |
| 2812836287 | MecB | 4 | 37.60 |
| 2812836288 | MecC | 6 | 36.88 |
| 2812836289 | MecD | 487 | 30.31 |
| 2812836290 | MecE | 15 | 36.15 |
| 2812836291 | MecF | 13 | 36.37 |
| 2812836292 | MecG | 1248 | 26.10 |
| 2812836293 | MecH | 68 | 34.00 |
| 2812836294 | MecI | 177 | 32.60 |

| '*Ca.* Dichloromethanomonas elyunquensis' | | | |
|---|---|---|---|
| **gene.ID** | **mec.protein** | **rank** | **log2.mean.abund** |
| 2628143003 | MecA2 | 607 | 21.81 |
| 2628142998 | MecB2 | 3 | 32.40 |
| 2628142997 | MecC2 | 4 | 32.15 |
| 2628142995 | MecE2 | 6 | 31.41 |
| 2628142994 | MecF2 | 9 | 30.57 |
| 2628142993 | MecG2 | 1669 | 16.09 |
| 2628142992 | MecH2 | 37 | 27.82 |
| 2628142991 | MecI2 | 83 | 26.42 |

**Table S2**. **Relative *mecE* transcript abundances in groundwater samples from a DCM-contaminated site.** Transcript relative abundance is expressed as transcripts per million transcripts (TPM).

| **id** | ***mecE* (TPM)** | **Location relative to DCM plume** |
|---|---|---|
| 47S104 | 2.69 | Core |
| GW47S | 8.43 | Core |
| WW01I | 21.24 | Core |
| GW01I100p | 0 | Core |
| GW01I99 | 4.39 | Core |
| 33I104 | 44.63 | Edge |
| 33I99p | 0 | Edge |
| 42SP | 0 | Edge |
| 48I104 | 0 | Outside |
| 48I99 | 0 | Outside |

**Table S3. Summary of *mec* cassette genes identified in metagenomes.** Metagenomes are grouped by general environmental source shown in column Genome.source; single.genome includes isolates and metagenome-assembled genomes (MAGs). For each *mec* gene ortholog identified, average amino acid identity (mean_ID) and total metagenomes, in which the homolog was identified (n), are provided.

| Genome.source | mec.ortholog | mean_ID | n |
|---|---|---|---|
| deep.subsurface | MecE | 63.99 | 2 |
| deep.subsurface | MecF | 66.69 | 2 |
| deep.subsurface | MecG | 40.61 | 2 |
| marine | MecB | 44.04 | 16 |
| marine | MecC | 37.97 | 17 |
| marine | MecE | 49.73 | 25 |
| marine | MecF | 51.58 | 18 |
| marine | MecG | 35.19 | 20 |
| marine | MecH | 39.44 | 24 |
| peat | MecA | 80.43 | 5 |
| peat | MecB | 72.03 | 10 |
| peat | MecC | 83.5 | 10 |
| peat | MecD | 82.66 | 8 |
| peat | MecE | 80.09 | 13 |
| peat | MecF | 90.87 | 8 |
| peat | MecG | 35.52 | 1 |
| peat | MecH | 35.3 | 1 |
| peat | MecI | 69.88 | 4 |
| single.genome | MecA | 87.89 | 7 |
| single.genome | MecB | 92.91 | 7 |
| single.genome | MecC | 94.25 | 7 |
| single.genome | MecD | 90.18 | 5 |
| single.genome | MecE | 94.13 | 5 |
| single.genome | MecF | 95.12 | 5 |
| single.genome | MecG | 88.78 | 4 |
| single.genome | MecH | 88.98 | 4 |
| single.genome | MecI | 91.15 | 5 |
| single.genome | MecJ | 70.58 | 6 |

**Table S4. Proximity of *mecJ* (encoding a putative cation antiport protein) homologs to dehalogenase-encoding genes.** The top IMG BLAST-P genome hits to *Dehalobacterium formicoaceticum* MecJ are shown. Adjacent dehalogenases were judged as being within 5 kb and in the same orientation as the MecJ homolog. Green shading indicates the presence of a dehalogenase-encoding gene: Reductive dehalogenase (RdhA, pfam13486) or S-2-haloacid dehalogenase (HADase, KEGG K01560). Orange shading indicates that the MecJ homolog is part of a putative anaerobic *mec* cassette described in this study. [2] indicates two of the designated genes that are present in immediate proximity. Gene ID is the IMG gene ID number.

| Gene ID | AA %ID | E-value | Bit Score | Organism | Adjacent dehalogenase |
|---|---|---|---|---|---|
| 2628143161 | 96.5% | 9.90E-266 | 827 | '*Ca.* Dichloromethanomonas elyunquensis' | *mec* cassette |
| 2628141546 | 77.6% | 2.50E-214 | 680 | '*Ca.* Dichloromethanomonas elyunquensis' | HADase [2] |
| 2753034167 | 63.5% | 2.80E-169 | 551 | *Dehalobacter* sp. FTH1 | RdhA |
| 2775644313 | 60.5% | 1.70E-158 | 521 | *Dehalobacter* sp. KB-1_124TCB1 | RdhA |
| 2578048931 | 57.5% | 2.80E-155 | 511 | *Dehalobacter* sp. UNSWDHB | RdhA [2] |
| 2520955686 | 57.5% | 2.80E-155 | 511 | *Dehalobacter* sp. 11DCA | RdhA [2] |
| 2520070062 | 57.5% | 2.80E-155 | 511 | *Dehalobacter* sp. CF | RdhA [2] |
| 2753034216 | 57.4% | 3.90E-153 | 505 | *Dehalobacter* sp. FTH1 | RdhA |
| 2578049036 | 53.1% | 1.10E-146 | 487 | *Dehalobacter* sp. UNSWDHB | *mec* cassette |
| 2776022298 | 52.9% | 7.60E-145 | 481 | Clostridiaceae bacterium mt12 | none |
| 2775643454 | 52.2% | 3.70E-143 | 476 | *Dehalobacter* sp. KB-1_124TCB1 | RdhA |

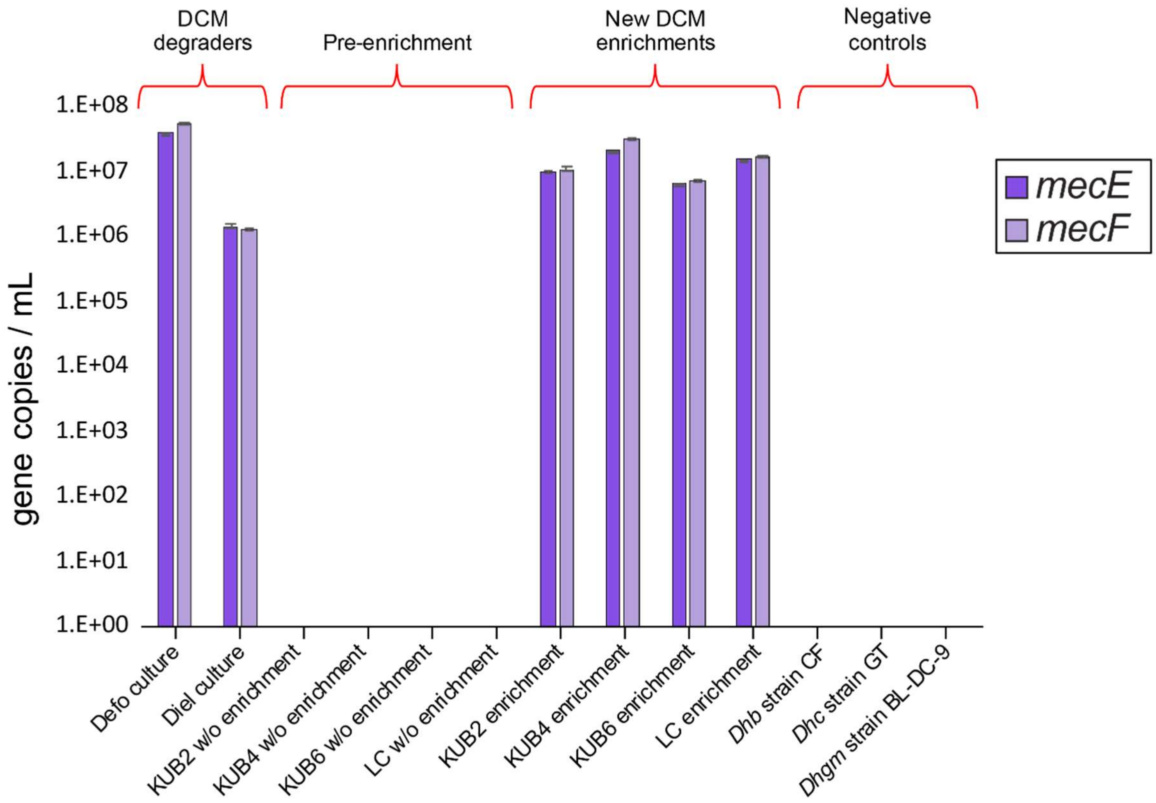


**Figure S1. Enrichment qPCR.** Results of *mecE* and *mecF* qPCR assays performed on anaerobic DCM enrichments derived from anaerobic municipal sewage sludge. DCM Degraders: *Dehalobacterium formicoaceticum* (Defo) and '*Ca*. Dichloromethanomonas elyunquensis' (Diel) present in enrichment culture RM grown with DCM; Pre-enrichment: sludge samples before enrichment on DCM; New DCM enrichments: anaerobic DCM enrichments from sewage sludges; Negative controls: assays performed on non DCM-degrading bacterial isolates; *Dhb*: *Dehalobacter* spp., *Dhgm*: *Dehalogenimonas* spp., *Dhc*: *Dehalococcoides mccartyi*.

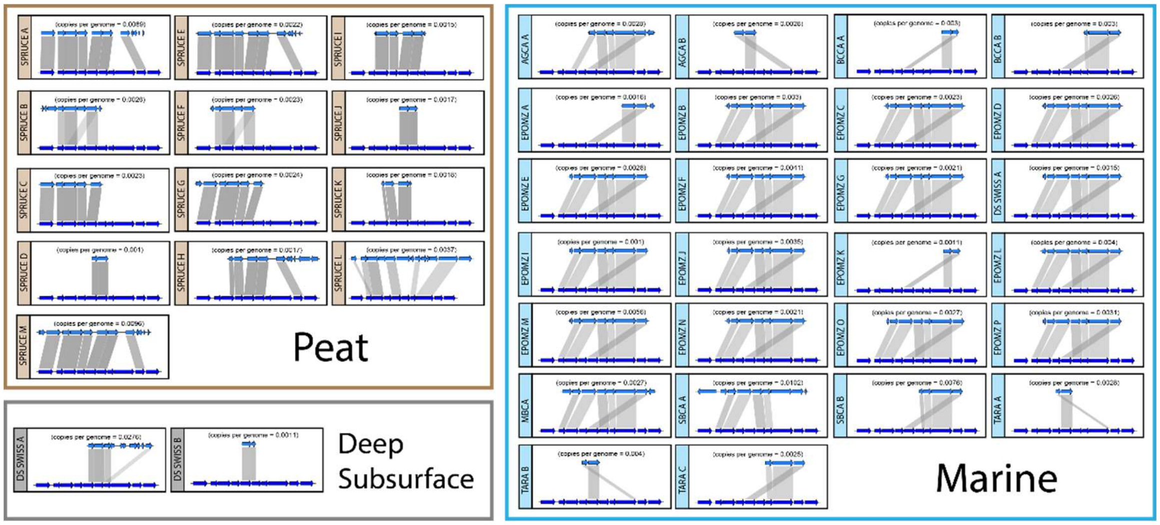


**Fig. S2**. **All *mec* cassette homologs identified in metagenomes available in the IMG database.** In each pairwise map, the Defo *mec* cassette is on the bottom and the metagenomic *mec* cassettes are on the top. The shades of grey are proportional to % amino acid identity. Specific IMG metagenome IDs for each cassette are listed in Dataset S7.

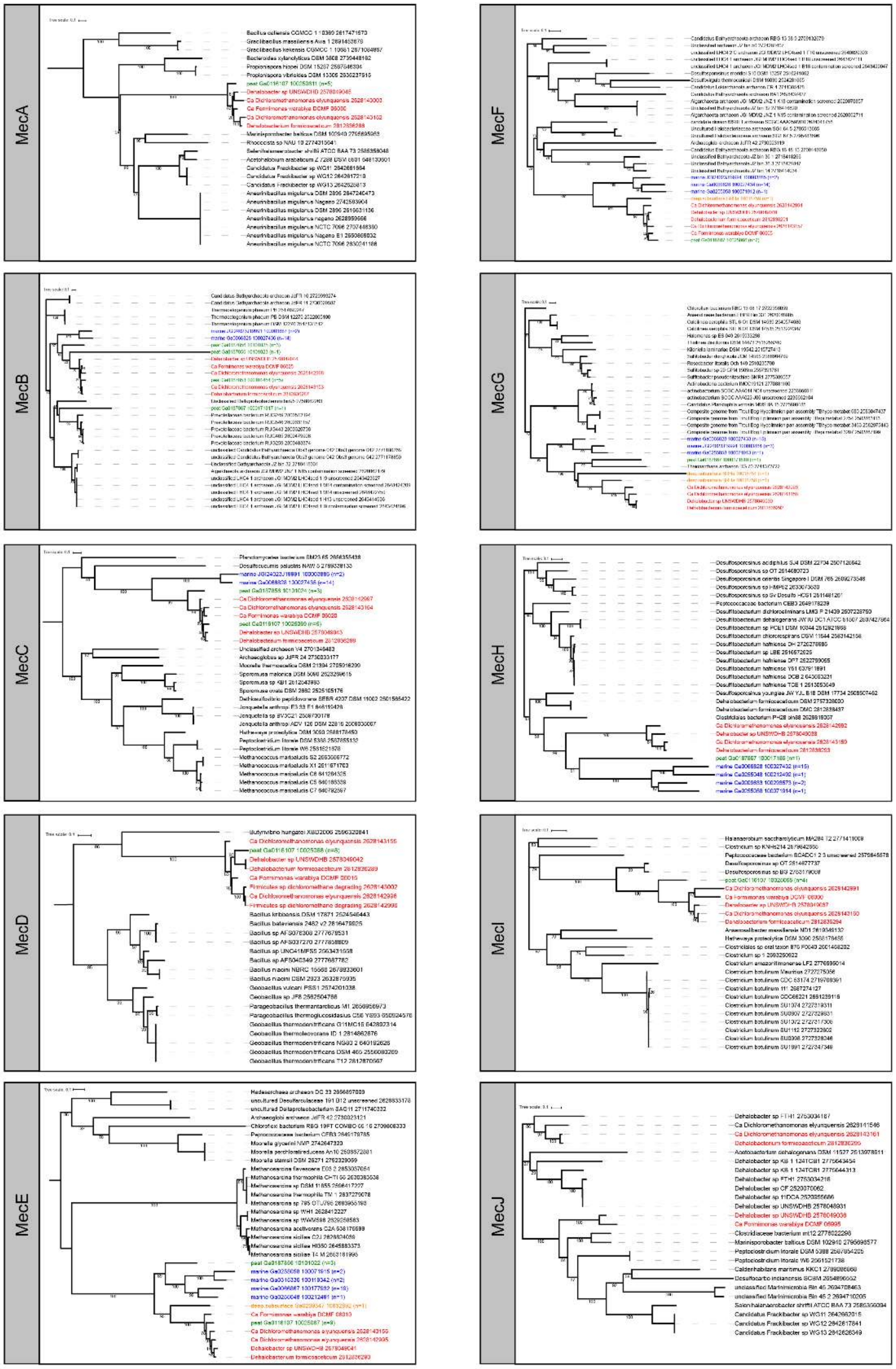
MecA
MecF
MecB
MecG
MecC
MecH
MecD
MecI
MecE
MecJ

**Figure S3.** Mec protein homologs encoded by the *mec* cassettes of Defo, Diel, Dcmf and metagenomes aligned and subjected to phylogenetic reconstruction alongside the top 20 most similar proteins encoded by microbial genomes in the IMG database. The top 20 homologs were identified based on BLAST-P similarity with a confidence cut-off of 1e-5. Leaf label color indicates the host genome source; Green indicates peat, blue indicates marine, orange indicates deep subsurface, and red indicates isolate or enrichment culture genome. Proteins encoded by genes located in metagenomes were clustered at 80% similarity using CD-Hit; the number of sequences each metagenome-derived leaf represents is indicated by n = x. Sequences were aligned using MAFFT G-INS-I with 1,000 maximum iterations and subjected to phylogenetic reconstruction using FastTree maximum-likelihood estimation (Gamma-LG model).

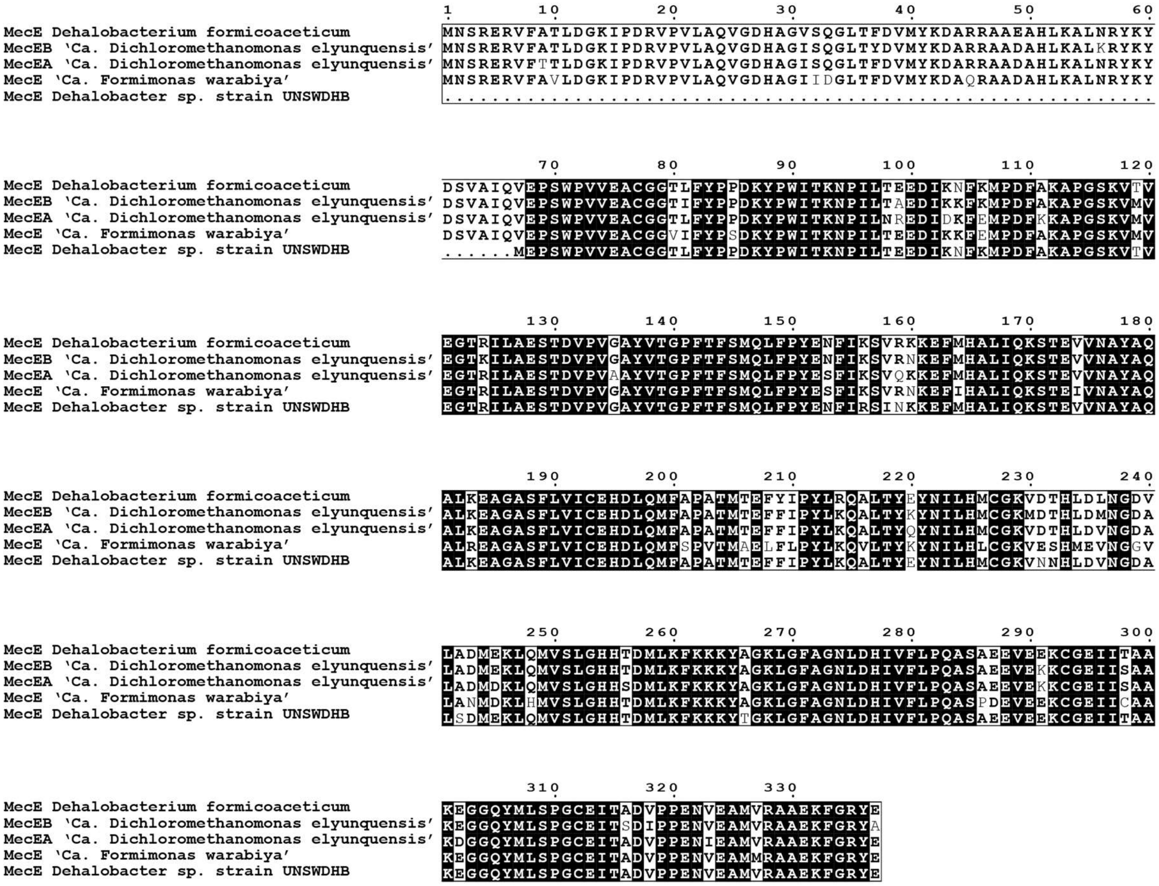


**Fig. S4. Alignment of all single-genome derived MecE homologs.** Sequences were aligned using MAFFT G-INS-I with 1,000 maximum iterations. Shaded residues represent full conservation across the alignment.

**Dataset S1 (separate file).** COG-defined single copy conserved protein (SCCP) read depths for genomes and metagenomes in which *mec* cassettes were identified. Average SCCP read depth was used to estimate total genome copies for use in calculating relative *mec* gene abundances in metagenomes (Dataset S7).

**Dataset S2 (separate file).** Full '*Ca*. Dichloromethanomonas elyunquensis' (Diel) proteome results. "RM.accession" indicates the culture RM IMG gene ID and Diel.accession indicates the IMG ID for the matching Diel gene. *mec* cassette homologs are indicated by the column labelled "MEC". $Log_2$ normalized abundances for three replicates and the averages are provided as indicated.

**Dataset S3 (separate file).** Sequences of primers, their amplification efficiencies, and standards used for qPCR studies.

**Dataset S4 (separate file).** Top 25 protein-level reciprocal best hits (RBH) between *Dehalobacterium formicoaceticum* and '*Ca*. Dichloromethanomonas elyunquensis'. RBH results are ranked in descending order of % amino acid identity. Calculations were performed using the IMG system.

**Dataset S5 (separate file).** Full *Dehalobacterium formicoaceticum* proteome results. "Gene.ID" indicates the IMG unique gene ID. *mec* cassette homologs are indicated by the column labelled "MEC". $Log_2$ normalized abundances for three replicates and the averages are provided as indicated.

**Dataset S6 (separate file).** Genome-derived orthologs to the mec cassette as anchored by homology searches based on *Dehalobacterium formicoaceticum* protein MecE. Requirements for MecE homology were minimum BLASTP bit-score of 150 and reciprocal best hit status.

Designation of surrounding genes comprising a *mec* cassette required the presence of at least one additional homolog using the same homology specifications. All identification numbers reference the IMG database except for genes from '*Ca*. Formimonas warabiya', where the numbers represent the NCBI codes. "MEC.cassette" is the codename given to the cassette as found in the indicated genome.

**Dataset S7 (separate file).** qPCR results for samples obtained from a DCM plume of a contaminated site and samples from two water columns of the Eastern Pacific Oxygen Minimum Zone (TNorth and St10). Average *mecE*, *mecF*, and bacterial 16S rRNA gene copy numbers (left) are provided along with standard deviations (far right). Average relative *mecE* or *mecF* to 16S rRNA gene abundances are provided in the center, along with the average ratio of *mecE* to *mecF* copy numbers.

**Dataset S8 (separate file).** All orthologs (genome- and metagenome-derived) to the putative DCM catabolism gene cassette as anchored by homology searches based on protein MecE of *Dehalobacterium formicoaceticum*. Requirements for MecE homology were a minimum BLAST-P bit-score of 150 and reciprocal best hit status. Designation of surrounding genes comprising a *mec* cassette required the presence of at least one additional homolog using the same homology specifications. All identification numbers reference the IMG database except for genes from '*Ca*. Formimonas warabiya', which represent the NCBI codes. "MEC.cassette" is the project-specific code name given to the cassette as found in the indicated genome or metagenome. "Genome.source" indicates the source of the (meta)genome, either the environment type (for metagenomes) or as a single genome, either an isolate or metagenome-assembled genome (MAG). "read.depth" is the copy number of the scaffold on which the cassette was located. "avg.sccp.read.depth" is the average copy number of 10 single-copy conserved protein-encoding genes (sccp) as defined by corresponding COG annotations as described in the Methods section. Full sccp counts for each (meta)genome are provided in Dataset S1. "genes.per.genome.copy" is cassette read depth divided by average sccp read depth, which provides an estimate of the number of genomes in the metagenome that contain the putative *mec* cassette homolog. "AA.ID" and e-value are the BLAST-P amino acid identity and e-value results of the indicated gene against the Defo-derived *mec* gene.

*Research*, *44*(D1), D372–D379. https://doi.org/10.1093/nar/gkv1103
